# ATLAS: Population-Level Disease Locus Discovery via Differential Attention in Genomic Language Models

**DOI:** 10.64898/2026.02.09.704696

**Authors:** Yuqi Liu, Kaiwen Deng, Yuhua Ye, Jiajie Zhan, Zilin Wang, Shicheng Chen, Xinyue Hu, An Chang, Zhaorong Li, Xin Jin, Shiping Liu, Kui Chen, Huijun Shen, Xianzhi Qi, Xiangmin Xu, Haiqiang Zhang

## Abstract

Identifying disease-associated genetic variants remains a key challenge in genomics, especially in small cohorts or for rare and complex mutation types where genome-wide association studies (GWAS) often fall short. We introduce ATLAS, a population-level framework that leverages attention signals from pretrained genomic language models (gLMs) to detect disease-associated genes and loci directly from raw sequences—without requiring explicit variant calls or supervised training. ATLAS first performs gene-level differential attention analysis to prioritize candidate genes, followed by base-level analysis to localize disease-associated regions at single-haplotype resolution. We validate ATLAS on synthetic and *β*-thalassemia datasets, demonstrating robust performance across diverse allele frequencies (down to 10%), cohort sizes (below 200 individuals per group), and genomic scales. Compared to GWAS, ATLAS achieves higher recall of known loci and captures haplotype-specific signals missed by traditional methods. Cross-model benchmarking further shows that precise localization depends on both model size and pretraining on diverse human genomes. In summary, ATLAS offers a scalable, sequence-native alternative to traditional statistical genetics.

## Introduction

Understanding how genetic variants affect gene activity is central to linking genotype to disease. Variants such as missense single-nucleotide polymorphisms (SNPs), insertions, and deletions can alter protein structure, disrupt regulatory elements, or perturb transcriptional regulation, contributing to Mendelian diseases, inherited metabolic disorders, and complex diseases including cancer and neurodevelopmental disorders Cutting (2015); Turner and Eichler (2019); Pagel et al. (2019); Montella et al. (2025). Accurately identifying such variants remains a central challenge in human genetics,with broad implications for disease prediction, mechanism discovery, and precision medicine.

Genome-wide association studies (GWAS) are the most widely used framework for identifying disease-associated variants, enabling the discovery of thousands of loci linked to complex traits through large-scale genotype–phenotype correlation analyses Risch and Merikangas (1996); Klein et al. (2005); Sollis et al. (2023). However, GWAS provide limited resolution at the haplotype level and are not optimized for non-SNP variants such as insertions and deletions Tewhey et al. (2011); Alkan et al. (2011). Their statistical power also depends strongly on allele frequency and effect size, often requiring very large cohorts to detect modest effects Visscher et al. (2017). Complementary bioinformatics tools predict variant effects using biological features such as sequence conservation and protein properties, avoiding large sample size requirements, but are typically restricted to predefined features and individual variants Adzhubei et al. (2013); Vaser et al. (2016).

Recent advances in genomic large language models (gLMs) provide a new paradigm for sequence-based variant analysis. Trained on massive genomic corpora, gLMs capture long-range dependencies and achieve state-of-the-art performance in regulatory annotation, variant effect prediction, and functional genomics DallaTorre et al. (2025); Brixi et al. (2025); Lin et al. (2025). Unlike GWAS, gLMs operate directly on raw DNA sequences and naturally accommodate heterogeneous variants, including substitutions and indels. However, most gLM-based applications focus on supervised prediction at the individual level Avsec et al. (2021); Benegas et al. (2023), leaving population-level sequence comparisons—analogous to GWAS but in a learned embedding space—largely unexplored.

In this paper, we propose **ATLAS** (Attention-based Locus Analysis System), an efficient attention-based explanation framework built on the genomic language model Genos Lin et al. (2025). We hypothesize that attention weights encode sequence-level importance and that functionally relevant loci manifest as statistically significant attention distribution shifts between case and control populations. By directly contrasting these attention patterns, ATLAS enables the identification of disease-associated loci without relying on explicit variant annotations or large sample sizes. Our main contributions are as follows.

- **End-to-end disease locus discovery**. We propose ATLAS, a framework that directly analyzes genomic sequences from multiple populations to identify disease-associated loci and genes without explicit variant calling.
- **Robust and flexible interpretation**. ATLAS is capable of multi-scale analysis at both gene and base levels with single-haplotype resolution, demonstrating robustness in small cohorts and high sensitivity to low-frequency variants.
- **Empirical validation on real-world data**. ATLAS not only recovers known disease-associated loci reported in the literature and GWAS but also identifies additional informative candidates on *β*thalassemia datasets.
- **Revisiting the value of human-centric foundation models**. By benchmarking diverse architectures, we demonstrate that massive parameter scale in generalist models does not guarantee performance. Instead, we conclude that accurate disease localization critically depends on the synergy between model capacity and extensive human-centric pretraining.

## Related Work

### Genomic Language Foundation Models

The application of large language models (LLMs) to genomics has shifted sequence analysis from alignment-based statistics to representation learning over raw DNA. These genomic language models (gLMs) are trained on large genomic corpora with self-supervised objectives to capture long-range dependencies and contextual sequence semantics. Representative general-purpose gLMs include LucaOne and the Nucleotide Transformer series, which leverage multispecies genomic data to learn transferable sequence representations Dalla-Torre et al. (2025); He et al. (2025). Evo 2 further extends this paradigm by modeling genomic sequences across all domains of life with strong generative capability Brixi et al. (2025). However, most general-purpose gLMs rely on reference genomes or cross-species consensus signals, limiting their sensitivity to human population variation and disease-specific sequence heterogeneity. Therefore, this work builds on Genos, a human-centric genomic language foundation model trained on large-scale human population sequencing data Lin et al. (2025), which has learned representations that are more sensitive to pathogenic variants and human-specific regulatory patterns.

### The Attention-based Model Interpretation

Attention mechanisms are intrinsic to Transformer architectures and provide an explicit, quantitative measure of how models weight long-range dependencies across input sequences. In genomics, attention has been widely used as an interpretability tool to reveal biologically meaningful interactions learned during supervised training. For example, Enformer demonstrated that attention maps capture long-range regulatory interactions underlying gene expression prediction Avsec et al. (2021). Beyond supervised tasks, attention has also been applied to unsupervised structural discovery, including the recovery of transcription factor binding motifs and the reconstruction of three-dimensional chromatin contact maps directly from sequences Tomaz da Silva et al. (2025); Boshar et al. (2025). While existing attention-based analyses predominantly focus on within-sequence interpretation for individual genomic sequences, our framework shifts attention analysis toward population-level comparison.

### Variant Effect Prediction Models

Prior work has largely focused on supervised variant effect prediction, estimating the impact of specific variant classes. Many approaches excel by specializing in defined biological mechanisms. For example, SpliceAI identifies splice-disrupting variants by modeling sequence context but is limited to splicing regulation Jaganathan et al. (2019). Similarly, AlphaMissense predicts missense pathogenicity using protein language models, yet remains restricted to coding regions Cheng et al. (2023). Unlike these variant-centric methods that evaluate mutations individually, ATLAS offers a model-agnostic pipeline capable of detecting diverse events without task-specific supervision or predefined variant classes.

### Statistical and Bioinformatic Approaches

Genome-wide association studies (GWAS) constitute the dominant statistical framework for genotypephenotype analysis, but their power critically depends on allele frequency, effect size, and large cohort sizes, requiring 10^4^–10^6^ samples to reliably detect variants at modest (10%) frequencies Visscher et al. (2017). Region-based aggregation methods such as SKAT partially alleviate this limitation by combining signals across predefined genomic regions or genes, yet still require moderate to large sample sizes for stable inference Wu et al. (2011); Zhan et al. (2016). In parallel, classical bioinformatic tools (e.g., SIFT4G, PolyPhen-2) estimate variant effects using curated annotations and prior biological knowledge, but are largely restricted to coding single-nucleotide substitutions and struggle to capture haplotype context or signals in poorly annotated regions Adzhubei et al. (2013); Vaser et al. (2016). Together, these limitations motivate alternative population-level frameworks that operate directly on sequence representations and are less dependent on explicit variant enumeration or extensive prior annotation.

## Methods

In this section, we present the ATLAS pipeline and its benchmarking against GWAS (Figure 1). We first describe how attention scores are extracted from genomic sequences using a pretrained genomic language model and how variable-length inputs are handled. We then introduce the gene-level differential attention analysis for identifying candidate genes. Next, we dive into these candidates, detecting disease-associated loci at baselevel resolution and clustering them into regions of interest. Finally, we summarize the GWAS procedures and describe how GWAS results are used for comparative evaluation. The code of ATLAS is available at https://github.com/BGI-HangzhouAI/ATLAS.

**Figure 1.**
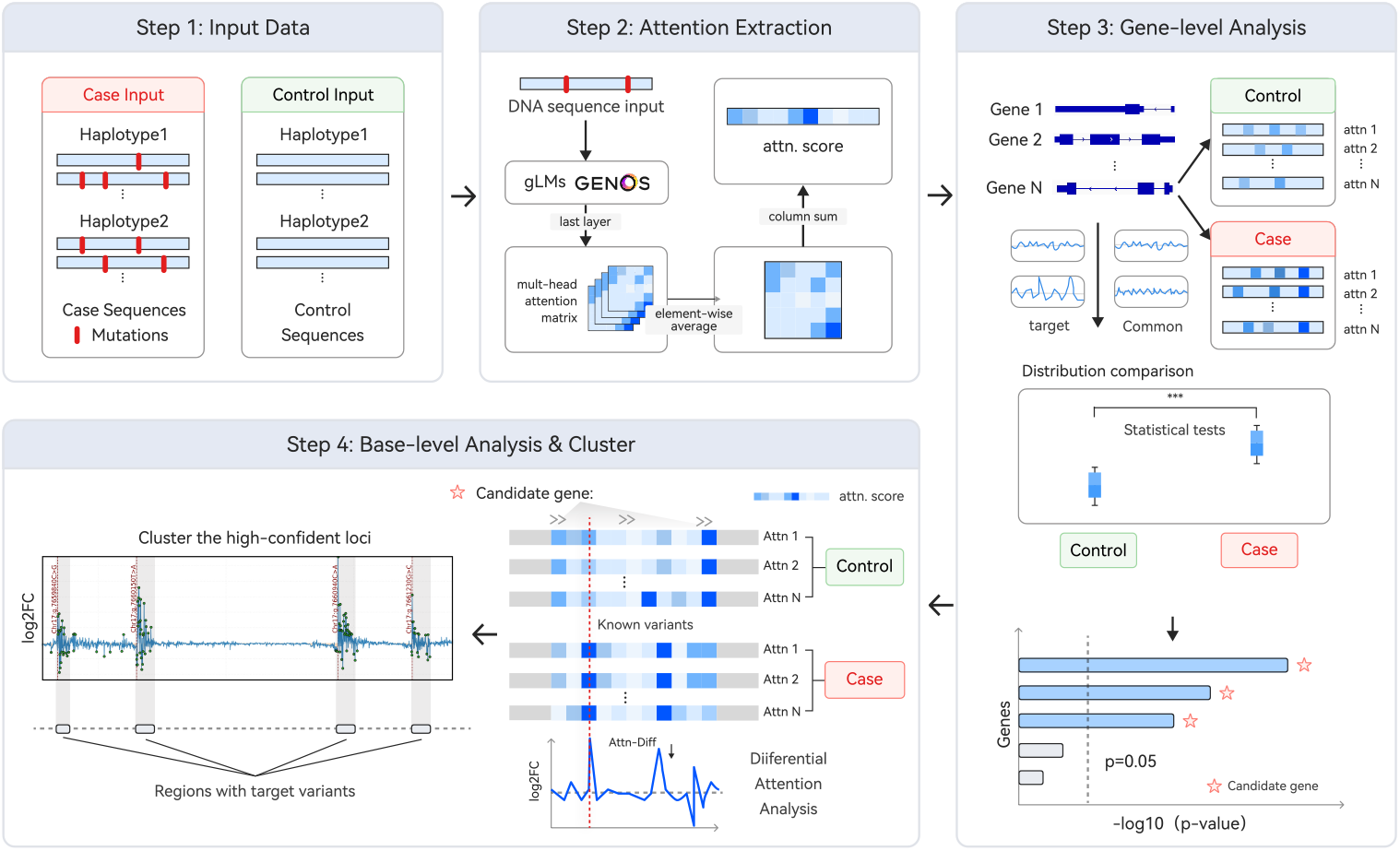
Overview of the ATLAS workflow. **Step 1:** ATLAS operates on haplotype-resolved genomic sequences derived from case and control cohorts. **Step 2:** Each sequence is processed a genomic language model (e.g., Genos). Multi-head attention matrices are extracted from the final layer, averaged element-wise, and aggregated via column-wise summation to obtain nucleotide-level importance scores. **Step 3:** ATLAS prioritizes candidate genes by performing statistical tests on attention score distributions between cohorts, identifying genes that exhibit significant disease-associated attention fluctuations relative to the background. **Step 4:** Within candidate genes, the framework calculates differential attention scores to identify specific loci with significant signal divergence. High-confidence sites are finally clustered to delineate discrete risk-associated genomic regions.

### Retrieve and Calculate Attention Scores

We extract attention scores from the final Transformer layer of each genomic language foundation model, with exceptions for specific architectures (e.g., the Evo 2 series Brixi et al. (2025); see Supplementary Note 3). This approach is motivated by evidence that the final layer captures the most globally contextualized sequence information Vig and Belinkov (2019) and serves as the primary interface for downstream biological tasks Marin et al. (2023). Consequently, we consistently employ final-layer attention to maximize the capture of highlevel semantic features.

Let the input sequence consist of *L* tokens. For each attention head *h*, we extract the query and key projections, **Q**^(*h*)^ and **K**^(*h*)^, and apply Rotary Position Embeddings (RoPE). We first define the pre-softmax attention matrix **A**^(*h*)^ incorporating the causal mask **M**_causal_. The final saliency score *s*_*j*_ for the *j*-th position is then derived by averaging the attention probabilities across *H* heads and summing the attention weights received from all query positions (i.e., column-wise summation):

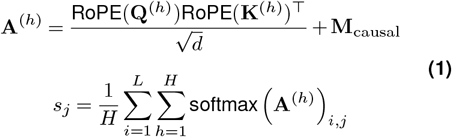

To efficiently compute attention scores across varying sequence lengths, we adopt a three-tiered strategy based on sequence length *L*. These thresholds are empirically determined to balance scoring latency and GPU memory consumptions on an NVIDIA H100 (80GB) GPU, ensuring numerical stability and signal integrity (see detailed benchmarks in Appendix C).

- **Vanilla attention (***L* ≤4, 096): For short sequences, we explicitly materialize the full *L*×*L* attention matrix. Our benchmarks indicate that within this range, the computational overhead is manageable, allowing full-matrix operations to be efficiently handled even on GPUs with more constrained memory capacities, such as the NVIDIA A40 (48GB).
- **FlashAttention-based (**4, 096 *< L*≤ 131, 072): For sequences up to 128k (where 1k = 1, 024), we employ a FlashAttention-style algorithm Dao et al. (2022). This approach computes *exact* attention scores using block-wise tiling and streaming softmax, effectively bypassing the *O*(*L*^2^) memory bottleneck while maintaining identical precision to vanilla attention.
- **Chunked processing (***L >* 131, 072): For even longer sequences exceeding the hardware’s single-pass capacity (128k), where memory consumption and computational costs grow quadratically, we apply a sliding-window chunking strategy. We empirically select a chunk size of 8, 192 with a 4, 096 overlap(Figure S1), as this configuration demonstrates the optimal balance between computational throughput and signal quality. To mitigate boundary effects and preserve continuity, we discard the outer half of each overlap region and retain only the central region of each chunk for final reconstruction.

### Gene-wise Differential Attention Analysis

We first identify candidate disease-associated genes using a gene-wise differential attention analysis, which compares attention distributions aggregated over defined genomic windows between control and disease cohorts. Prior work has shown that sequence variants can induce systematic and biologically meaningful perturbations in attention patterns of genomic Transformer models, with detectable fluctuations in aggregated attention distributions Consens et al. (2025). Motivated by this evidence, we hypothesize that genes involved in disease will exhibit reproducible differences in attention distributions relative to controls.

For each gene, we consider the entire gene body. The start and end positions are retrieved directly from the Ensembl database. We summarize the per-base attention scores across these regions for each sample using a set of complementary statistics designed to capture central tendency, extremum, and dispersion (Supplementary Note 5). For each summary statistic, we compare the distributions between control and disease cohorts using a two-sided Mann–Whitney U test. To account for multiple comparisons across the genome, we apply the Benjamini-Hochberg procedure to control the False Discovery Rate (FDR). Genes with FDR-adjusted *p*-values *<* 0.05 are considered significant.

Across our evaluations, Shannon Entropy consistently exhibits the strongest discrimination. We attribute this to the model’s sensitivity to risk syntax: rather than dispersing attention, disease-associated variants act as strong attractors. This causes the model’s focus to systematically concentrate on specific loci, thereby reducing the distributional entropy compared to controls. Consequently, Shannon Entropy is emphasized in subsequent analyses.

### Base-wise Differential Attention Analysis

To localize potentially disease-associated loci within the candidate genes prioritized in Section, we perform a base-level differential attention analysis. Drawing an analogy to differential expression analysis used to identify disease signatures Rosati et al. (2024), we hypothesize that high-risk loci induce systematic fluctuations in attention patterns between case and control cohorts.

For each genomic position *j*, we aggregate attention scores across valid samples in each cohort. Let 𝒮_*c*_ and𝒮_*d*_ denote the sets of samples where position *j* is effectively present (i.e., non-deleted and non-padding) in the control and disease groups, respectively. To rigorously handle Indels, we align comparisons to the reference coordinates: for deletions (e.g., AAAT →A), differential attention is quantified at the retained anchor (start) position. Positions falling within the deleted span are excluded from 𝒮_*g*_ for the affected samples to prevent zero-inflation artifacts. We compute the base-wise log_2_ fold change (LFC_*j*_) as:

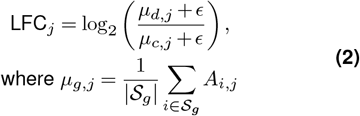

where *A*_*i,j*_ represents the attention score of sample *i* at position *j*, and *ϵ* is a small pseudo-count to ensure numerical stability.

The statistical significance of this difference is assessed using a two-sided Mann–Whitney U test, with *p*-values adjusted via the Benjamini–Hochberg (BH) procedure. A position is identified as a *highly differentiated attention locus* if it satisfies two criteria: (1) adjusted *p*-value *<* 0.01; and (2) absolute LFC_*j*_ exceeds a length-adaptive quantile threshold (*q*_*L*_). This dynamic thresholding strategy is critical for maintaining a constant false discovery rate across varying genomic contexts, as detailed in Section .

### Clustering and Delineating Disease-associated Regions

We observe that differential attention signals do not always strictly co-localize with target variants at singlebase resolution; rather, they exhibit a proximal enrichment pattern, where significant attention fluctuations spatially cluster around disease-associated loci. Consequently, we aim to aggregate these discrete highconfidence signals into contiguous candidate risk regions.

#### Adaptive Thresholding

To ensure consistent detection sensitivity, the filtering threshold *q*_*L*_ introduced in Section is formulated as a function of sequence length *L*. We empirically calibrate the baseline threshold on 4 kb sequences, identifying the 95th percentile (*α* = 0.05) as the optimal cutoff for effective signal isolation. However, applying this fixed threshold to longer sequences results in elevated false discovery rates due to the expanded search space. To counteract this, we define *q*_*L*_ (for *L* in kb) as:

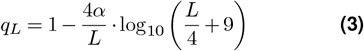

This formulation is strictly grounded in our experimental observations: the term is designed to satisfy the boundary condition *q*_*L*_ = 1 −*α* = 0.95 at the baseline length (*L* = 4 kb), while imposing progressively stricter filtering constraints (asymptotically approaching 1) for longer sequences to rigorously suppress the accumulation of background noise.

#### Density-based Clustering

We apply DBSCAN Ester et al. (1996); Schubert et al. (2017) to the loci satisfying the *q*_*L*_ criterion, calculating clusters along 1D genomic coordinates using Euclidean distance. We set the neighborhood radius *ϵ* = 20 bp, approximating the typical length of transcription factor binding sites or regulatory motifs, and min_samples = 5 to ensure that detected clusters represent robust, multi-base signal accumulations rather than isolated artifacts.

### Benchmark with GWAS

To establish a baseline comparison with traditional statistical methods, we performed genome-wide association studies (GWAS) on genotype data from 1,429 *β*-thalassemia carriers and patients (details in Supplementary Note 2 A).

We implemented a stringent quality control (QC) pipeline using VCFtools Danecek et al. (2011). First, multi-allelic variants were split into biallelic records, and chromosome identifiers were standardized. We then applied genotype-level filtering to set genotypes with sequencing depth (DP) *<* 4 to missing. Subsequently, variant-level filtering was performed to retain only autosomal loci with a missingness rate *<* 15% and a minor allele frequency (MAF) ≥1%.

The post-QC dataset was converted to PLINK binary format using plink2 Chang et al. (2015). We performed a case–control association analysis using a logistic regression model to test the additive effect of each variant on the binary disease phenotype. To control for false positives, statistical significance was assessed using the conventional genome-wide significance threshold (*p <* 6.3 × 10^−9^).

### Experiment Design

#### Synthetic Datasets

To systematically evaluate ATLAS’s robustness under controlled conditions—particularly for small cohorts and low allele frequencies—we constructed a series of synthetic datasets using background noise derived from the real *β*-thalassemia cohort (details in Supplementary Note 2 A).

We sampled four genomic regions of increasing length (4 kb, 20 kb, 128 kb, and 384 kb) from the reference genome, explicitly excluding the *HBB* locus to prevent data leakage. Within each region, synthetic target variants were inserted as ground truth, simulating increasing complexity: from 8 SNPs in 4 kb regions to a mixture of 20 variants (15 SNPs + 5 Indels) in larger windows. Genotypes were assigned based on disease status: cases were modeled as homozygous mutants (1| 1) and controls retained the reference genotype (0 |0).

We designed two experimental settings to stress-test the model:

- **Allele Frequency (AF) Sensitivity:** In the 4 kb region, we simulated lower penetrance by randomly downsampling mutant genotypes (1 |1 →0| 0) in case samples to target frequencies of 70%, 50%, 20%, and 10%, while keeping background sites unchanged.
- **Sample Efficiency:** We varied the balanced case–control cohort sizes from 200:200 down to 10:10, fixing the penetrance of synthetic target variant at 100%.

Sequence construction followed a strict coordinate mapping pipeline to handle Indel-induced shifts (see Supplementary Note 2 B).

### Thalassemia Cohort and Data Preprocessing

We designed this real-world evaluation to assess whether ATLAS can recover known risk loci under realistic population heterogeneity and, crucially, to identify plausible disease-associated signals beyond curated annotations.

The *β*-thalassemia dataset was derived from a crosssectional whole-genome sequencing study investigating the clinical heterogeneity of hemoglobinopathies comprising 1,429 individuals. For validation, we referenced the IthaGenes database Kountouris et al. (2014), which documents 512 thalassemia-associated variants in the *HBB* gene. Within our cohort, 31 of these recorded variants were identified. We selected the 8 most prevalent variants routinely used in clinical diagnosis as the ground-truth set for our primary performance benchmarks Writing Group For Practice Guidelines For Diagnosis And Treatment Of Genetic Diseases Medical Genetics Branch Of Chinese Medical Association et al. (2020).

Raw genotypes were processed to generate model-ready sequences. First, VCF records were phased using Beagle4 Browning and Browning (2007) to obtain haplotype-resolved genotypes. Subsequently, we reconstructed full haplotype sequences for each individual by applying variants to the GRCh38 reference genome. Specific rules for handling Indels, coordinate mapping, and reverse-complementation for negative-strand genes are detailed in Supplementary Note 2 B.

#### Genome-Wide Scanning on Chromosome 11

To evaluate beyond known *HBB* mutations, we extended the analysis to all protein-coding genes on chromosome 11. This genome-wide scan aimed to: (i) identify previously uncharacterized loci; and (ii) evaluate method specificity, under the expectation that non–hematology-related genes should exhibit minimal differential attention signals.

### Evaluation Metric

#### Base-Resolution Localization Accuracy

To evaluate localization performance at single-base resolution, we measure the extent to which attention-derived signals concentrate near known disease-associated variant positions. We posit that informative signals should preferentially localize in the immediate vicinity of causal variants; distal signals are considered less actionable for downstream validation.

Given the extreme class imbalance between causal variants (rare) and background bases (abundant), we adopt the Area Under the Precision–Recall Curve (AUPRC) as our primary metric. We utilize absolute log_2_ fold-change (log_2_ FC) values as scores, treating bases within fixed windows around each variant as positive labels. To provide a comprehensive assessment of signal quality beyond ranking, we report three complementary metrics:

- **Signal-to-Noise Ratio (SNR):** Measuring the magnitude contrast between signal and background regions.
- **Fraction of Signal in Windows (FRiW):** Quantifying the efficiency of attention mass allocation (analogous to ChIP-seq FRiP scores).
- **Weighted Distance (WDist):** Assessing spatial precision without imposing hard window boundaries.

For cross-scale comparisons (e.g., 4 kb vs. 384 kb sequences), we utilize length-normalized variants of FRiW and WDist, alongside the mean percentile rank of true positions. Detailed mathematical definitions are provided in Supplementary Note 4.

#### Cluster-Level Recovery

We further evaluate performance at the region level by assessing whether the contiguous clusters identified by our pipeline (Section) successfully overlap with known variant loci. For synthetic datasets, we compute both precision and recall to measure the recovery of predefined variants and the suppression of spurious clusters. For real-world cohorts (e.g., *β*-thalassemia), where ground-truth annotations are inherently incomplete, we prioritize recall—quantifying the coverage of clinically validated loci. Clusters lacking overlap with annotated variants are not strictly penalized as false positives, as they may represent novel, uncharacterized biological signals.

## Results

In this section, we present the evaluation of ATLAS in three parts. First, we demonstrate its practical efficacy by identifying disease-associated genes and localizing fine-grained signals in a real-world *β*-thalassemia cohort. Second, we validate the method’s robustness under controlled conditions using synthetic datasets with varying cohort sizes and signal strengths. Finally, we benchmark different foundation models to reveal how model scale and data diversity impact disease localization performance.

### Gene-level differential attention identifies disease-associated genes

Our gene-level differential attention analysis successfully prioritizes the *HBB* gene as the top candidate on chromosome 11, demonstrating the model’s capacity to distinguish disease-associated signals from the genomic background. As illustrated in Figure 2, *HBB* dominates the rankings across both haplotypes, achieving significance levels (−log_10_ *p*_adj_) that are 6.36 ×and 9.60 ×higher than the second-ranked candidates on haplotype 1 and haplotype 2, respectively. Specifically, disease samples exhibite a sharp reduction in Shannon entropy within the *HBB* locus compared to controls (*p*_adj_ *<* 10^−32^, Figure 3). This significant drop indicates that the model’s attention—typically dispersed in controls—becomes systematically focused on specific disease-associated sites in the disease state. This pattern proves robust across multiple distributional metrics, including standard deviation and coefficient of variation (Figure S5 and S6).

**Figure 2.**
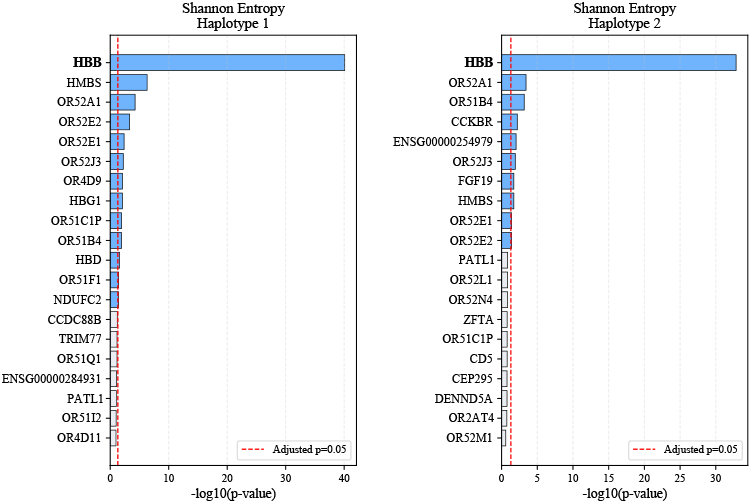
Top-20 genes with differently distributed attentions quantified in entropy. The bar plots rank the top-20 protein-coding genes on chromosome 11 by −log_10_(*p*) values from Wilcoxon rank-sum tests for Shannon entropy, highlighting HBB as the most significant gene.

**Figure 3.**
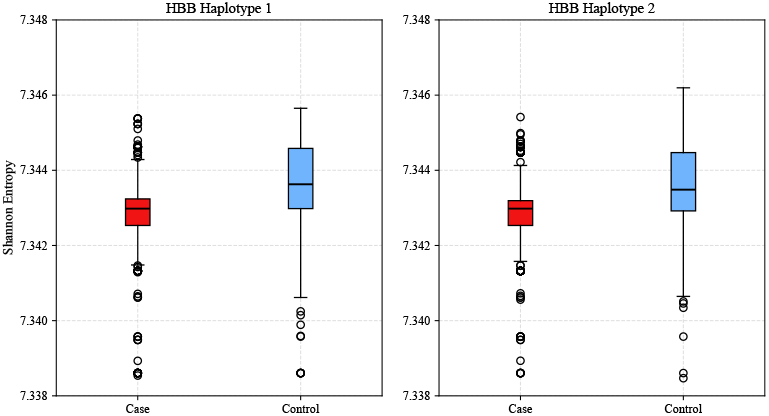
Comparison of Shannon entropy values of HBB in case and control groups. Both haplotypes show significantly lower entropy values in the case group compared to the control group.

Beyond the expected *HBB* signal, the analysis demonstrates high specificity. Among the top candidates, only 15 other genes surpass the significance threshold (*p*_adj_ *<* 0.05) across the entire chromosome. Crucially, four of these genes—*HBG1, HBD, HBMS*, and *OR52A1*—are biologically validated as thalassemia modifiers or hemoglobin regulators (Table 1). The recovery of these secondary but functionally relevant genes, amidst a low false-positive background, confirms that differential attention entropy serves as an effective, zero-shot filter for isolating disease-relevant loci prior to fine-grained mapping.

**Table 1.** Biological evidence for top-ranked *β*-thalassemia associated genes identified by ATLAS.

| Gene | Biological Relevance |
| --- | --- |
| HBMS | β-thalassemia associated gene inferred by GeneCard Stelzer et al. (2016) and DISEASE databases Grissa et al. (2022). |
| OR52A1 | Contains the γ-globin enhancer; variants in this region can affect erythropoiesis and modulate thalassemia phenotypes Himadewi et al. (2021). |
| HBG1 | A key modifier gene encoding γ-globin. Up-regulation (↑HbF) ameliorates clinical severity in β-thalassemia. |
| HBD | Encodes δ-globin. Co-inherited variants or altered HbA<sub>2</sub> levels are diagnostic hallmarks for β-thalassemia carriers. |

### Synthetic validation of base-level localization accuracy

We evaluate ATLAS on synthetic datasets to quantify robustness under varying conditions, reporting metrics averaged across window sizes and haplotypes. The 4 kb dataset with 100% allele frequency and 1,429 samples serves as the primary baseline.

ATLAS demonstrates high resilience to data sparsity and rare variants (Table 2, Figure S3). Notably, clear signal concentration around synthetic target sites remained observable even at 10% allele frequency or with as few as 10 individuals per group. Base-level metrics (AUPRC, SNR, FRiW) show minimal degradation (*<* 10%) under these challenging conditions, while cluster-level precision and recall consistently remained above 0.80. This confirms that ATLAS can reliably prioritize rare risk variants even in small-scale studies.

**Table 2.**
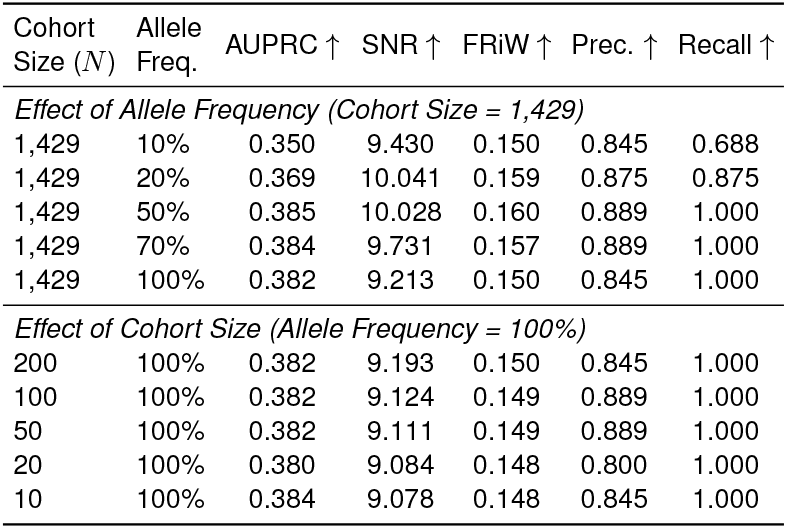
Base-level localization and cluster-level recovery performance on 4 kb synthetic sequences under varying cohort sizes and allele frequencies.

| Cohort Size ($N$) | Allele Freq. | AUPRC ↑ | SNR ↑ | FRIW ↑ | Prec. ↑ | Recall ↑ |
| --- | --- | --- | --- | --- | --- | --- |
| *Effect of Allele Frequency (Cohort Size = 1,429)* |  |  |  |  |  |  |
| 1,429 | 10% | 0.350 | 9.430 | 0.150 | 0.845 | 0.688 |
| 1,429 | 20% | 0.369 | 10.041 | 0.159 | 0.875 | 0.875 |
| 1,429 | 50% | 0.385 | 10.028 | 0.160 | 0.889 | 1.000 |
| 1,429 | 70% | 0.384 | 9.731 | 0.157 | 0.889 | 1.000 |
| 1,429 | 100% | 0.382 | 9.213 | 0.150 | 0.845 | 1.000 |
| *Effect of Cohort Size (Allele Frequency = 100%)* |  |  |  |  |  |  |
| 200 | 100% | 0.382 | 9.193 | 0.150 | 0.845 | 1.000 |
| 100 | 100% | 0.382 | 9.124 | 0.149 | 0.889 | 1.000 |
| 50 | 100% | 0.382 | 9.111 | 0.149 | 0.889 | 1.000 |
| 20 | 100% | 0.380 | 9.084 | 0.148 | 0.800 | 1.000 |
| 10 | 100% | 0.384 | 9.078 | 0.148 | 0.845 | 1.000 |

Signal localization scales effectively across genomic contexts. As shown in Table 3, we extend the evaluation to sequences ranging from 4 kb to 384 kb. Length-normalized metrics demonstrate consistent performance: *FRiW Enrichment* scales with sequence length, indicating effective signal isolation against expanding genomic backgrounds, while *Normalized Weighted Distance* remained consistently low (*<* 0.4). Notably, for the longest sequences (384 kb), cluster-level recovery achieves near-perfect scores (Precision/Recall ≈1.0), validating the pipeline’s capability to detect sparse signals in large genomic windows (Figure S4).

**Table 3.** Performance scaling across increasing sequence lengths (4 kb to 384 kb).

| Seq. Length | FRIW Enrich. ↑ | Normalized WDist ↓ | Mean Rank ↑ | Prec. ↑ | Recall ↑ |
| --- | --- | --- | --- | --- | --- |
| 4 kb | 8.03 | 0.373 | 0.926 | 0.844 | 1.000 |
| 20 kb | 19.04 | 0.328 | 0.908 | 0.909 | 1.000 |
| 128 kb | 63.41 | 0.252 | 0.908 | 0.950 | 0.950 |
| 384 kb | 129.82 | 0.325 | 0.915 | 1.000 | 1.000 |

### Base-level discovery of disease-associated regions

We next apply base-level differential attention analysis to top candidate genes, where ATLAS demonstrates superior recall of known disease-associated variants in *HBB* compared to standard GWAS. We compare the detected attention clusters against both GWAS results (*p <* 6.3 ×10^−9^) and the 31 clinical variants reported in IthaGenes. As illustrated in Figure 4, while GWAS identify only a single locus overlapping with known variants, ATLAS detects 4 distinct clusters on haplotype 1 (covering 7 reported sites) and 3 clusters on haplotype 2 (covering 6 reported sites). This confirms that attention-derived signals can recover a substantially larger proportion of clinically reported variants that are statistically elusive to standard association mapping.

**Figure 4.**
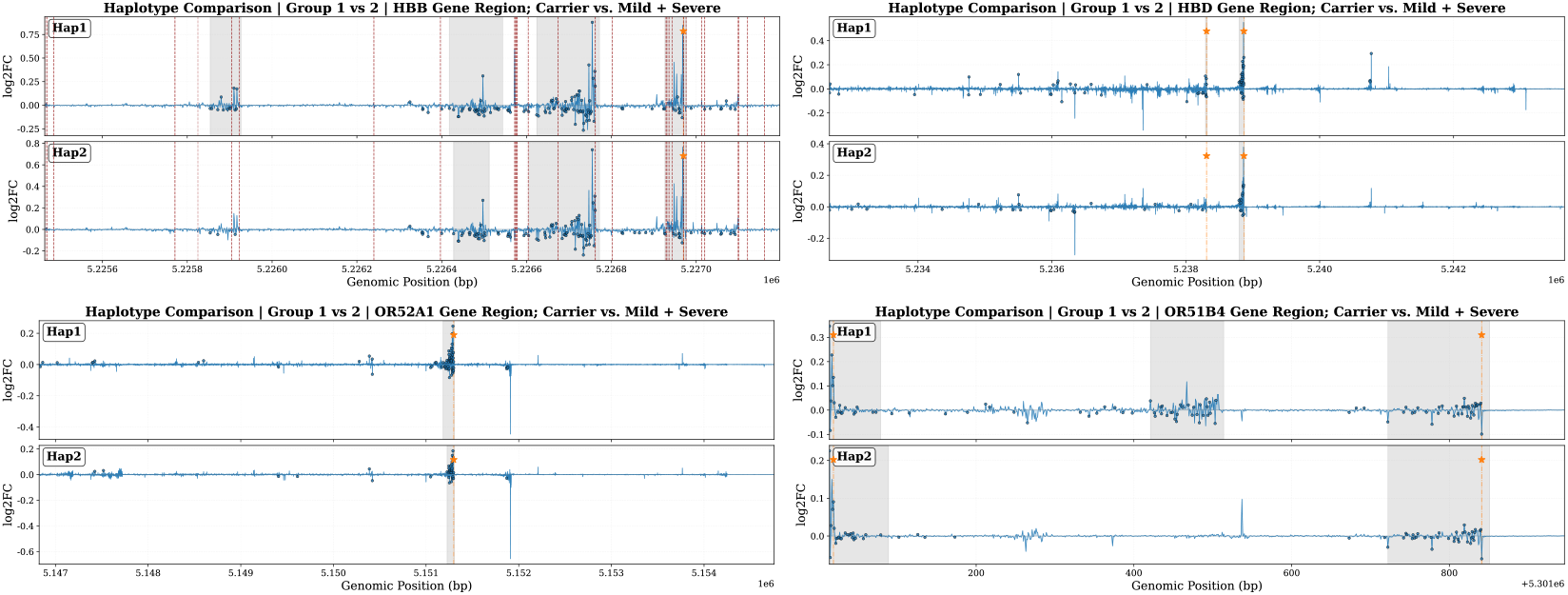
Comparision of the derived cluster, known loci, and GWAS inference on both haplotypes. Here we visualize the four genes that both GWAS and ATLAS have found. The gray regions are the clusters with their cluster score showed as the colored dots. The red lines are the known loci. The orange star is the GWAS inferred locus.

Extending the analysis to the other 15 candidate genes, ATLAS shows strong agreement with standard methods while offering superior spatial resolution. Our method successfully recovers all GWAS-significant loci (e.g., in *OR52A1, HBD*) and identifies latent signals in 6 additional genes. Notably, the detected clusters span an average of only 11.62 bp. This fine-grained localization significantly narrows down the search space for putative risk variants, providing much tighter candidate windows compared to typical LD-based association blocks. Furthermore, ATLAS leverages haplotype information to reveal phase-specific patterns—such as the asymmetric attention clusters in *HBD*—which are typically obscured in population-based GWAS signals.

### Impact of model capacity and pretraining diversity on loci discovery

Furthermore, we explore how model architecture and pretraining data influence the localization of human disease variants. We benchmark models with varying scales and training domains—including the Evo2 series, Luca series, and Genos family—on the *β*-thalassemia cohort (Table 4).

**Table 4.** Foundation-model comparison on *β*-thalassemia (HBB) under ATLAS.

| Model | Pretrain domain | Avg. AUC ↑ | Avg. AUPRC ↑ | Avg. SNR ↑ | Avg. FRIW ↑ |
| --- | --- | --- | --- | --- | --- |
| Genos-v2 | Human (diverse) | 0.807 | 0.265 | 4.941 | 0.169 |
| Genos-v1 | Human | 0.804 | 0.258 | 4.496 | 0.157 |
| Genos-1.2b | Human | 0.777 | 0.173 | 3.237 | 0.121 |
| Evo2-7b | Generalist | 0.712 | 0.147 | 2.566 | 0.096 |
| Evo2-1b | Generalist | 0.753 | 0.141 | 2.782 | 0.107 |
| Evo2-40b | Generalist | 0.709 | 0.093 | 2.139 | 0.082 |
| LucaOne | Generalist | 0.687 | 0.094 | 2.148 | 0.081 |
| LucaVirus | Virus | 0.695 | 0.087 | 1.908 | 0.074 |

First, our observations indicate that model capacity is a significant factor in signal quality, particularly when the domain aligns. Within the human-centric Genos family, scaling up parameters yielded consistently cleaner signals. As shown in Table 4, Genos-v2 substantially improves metrics over Genos-1.2B (AUPRC: 0.173→0.265; SNR: 3.24 →4.94), suggesting that larger models benefit from enhanced semantic denoising capabilities in non-informative regions.

Second, the results also suggest that scaling parameters alone may not guarantee performance gains if the pretraining data is not sufficiently aligned with the target task. This is notable in the comparison with the generalist Evo2-40B model: despite its substantial parameter count, it does not exhibit a commensurate performance advantage in localizing human pathogenic variants (Avg AUPRC ≈0.093), performing similarly to smaller baselines. In contrast, Genos-v2, which benefits from extensive human genome diversity, consistently outperforms generalist counterparts.

The findings imply that for specific human disease tasks, achieving reliable localization performance likely requires a synergy between sufficient model capacity and diverse human-centric pretraining.

## Discussions

### Conclusion

We present ATLAS, a framework that establishes a new paradigm for population-level discovery through the lens of genomic language models. By decoding internal attention patterns, ATLAS bypasses the limitations of traditional frequency-based statistics, enabling the discovery of disease-associated loci directly from sequence semantics.

Our results confirm that identifying risk variants does not strictly require massive cohorts or clear statistical separation; rather, actionable signals can be extracted from the model’s intrinsic understanding of genomic syntax. ATLAS thus positions itself not merely as a complement to GWAS, but as a critical instrument for illuminating the “genetic dark matter”—including rare variants, complex haplotypes, and non-coding regions—that remains inaccessible to conventional methods. Ultimately, this work paves the way for a future where genomic language models become the standard engine for decoding the complex syntax of human disease.

### Limitations and Future Work

First, this study focuses on binary phenotypes driven by major genes; extending ATLAS to complex, polygenic traits remains a key future direction. Second, our analysis currently relies on reconstructed genomes with inherent processing noise. As high-fidelity sequencing data becomes more accessible, ATLAS is expected to leverage these improved inputs to naturally enhance localization precision without architectural changes. Finally, while ATLAS effectively prioritizes candidate loci, establishing definitive causality warrants further empirical verification. Future work will focus on validating these computational predictions through large-scale functional assays.

## Ethics Statement

The *β*-thalassemia dataset utilized in our research are from the study approved by the ethics committee of Nanfang Hospital, Southern Medical University, and the ethical committees of each local hospital participating in this study (Approval No. NFEC-2019-039). All subjects and/or their guardians provided written informed consent. The data supporting the findings of this study will be made available upon reasonable request to the corresponding author, subject to approval by the original data custodian(s).

## Supplementary Note 1: Implementation Details & Reproducibility

To ensure the reproducibility of our experiments and the scalability of our approach to long-context genomic sequences, we provide detailed specifications of our computational environment, attention extraction algorithms, and hyperparameter optimization strategies.

### A. Computational Environment

All models were trained and evaluated on a high-performance computing node optimized for large-scale deep learning.

- **Hardware Infrastructure:** A single node equipped with 4× NVIDIA A40 GPUs (48GB VRAM per GPU). This setup utilizes the Ampere architecture to support efficient `bfloat16` mixed-precision training.
- **Software Stack:**
  – **Framework:** PyTorch `2.5.1+cu124` with Python 3.12.2.
  – **Acceleration: FlashAttention-2** (v2.8.3) was employed to optimize the memory hierarchy (HBM vs.SRAM), significantly reducing the IO overhead for attention matrix computation.
  – **CUDA Toolkit:** Version `12.4.`

### B. Attention Score Extraction & Alignment

Unlike standard NLP tasks where token indices align linearly with positions, genomic analysis requires precise mapping between attention weights and biological coordinates, especially in the presence of Indels.

#### B.1 FlashAttention-based Extraction Logic

We extract raw per-token attention importance scores directly from the Transformer layers. To handle the quadratic complexity of attention in long sequences, we implement a block-wise extraction algorithm based on FlashAttention logic. The detailed procedure is formally described in Algorithm 1.

##### Algorithm 11

FlashAttention-based Per-token Attention Extraction

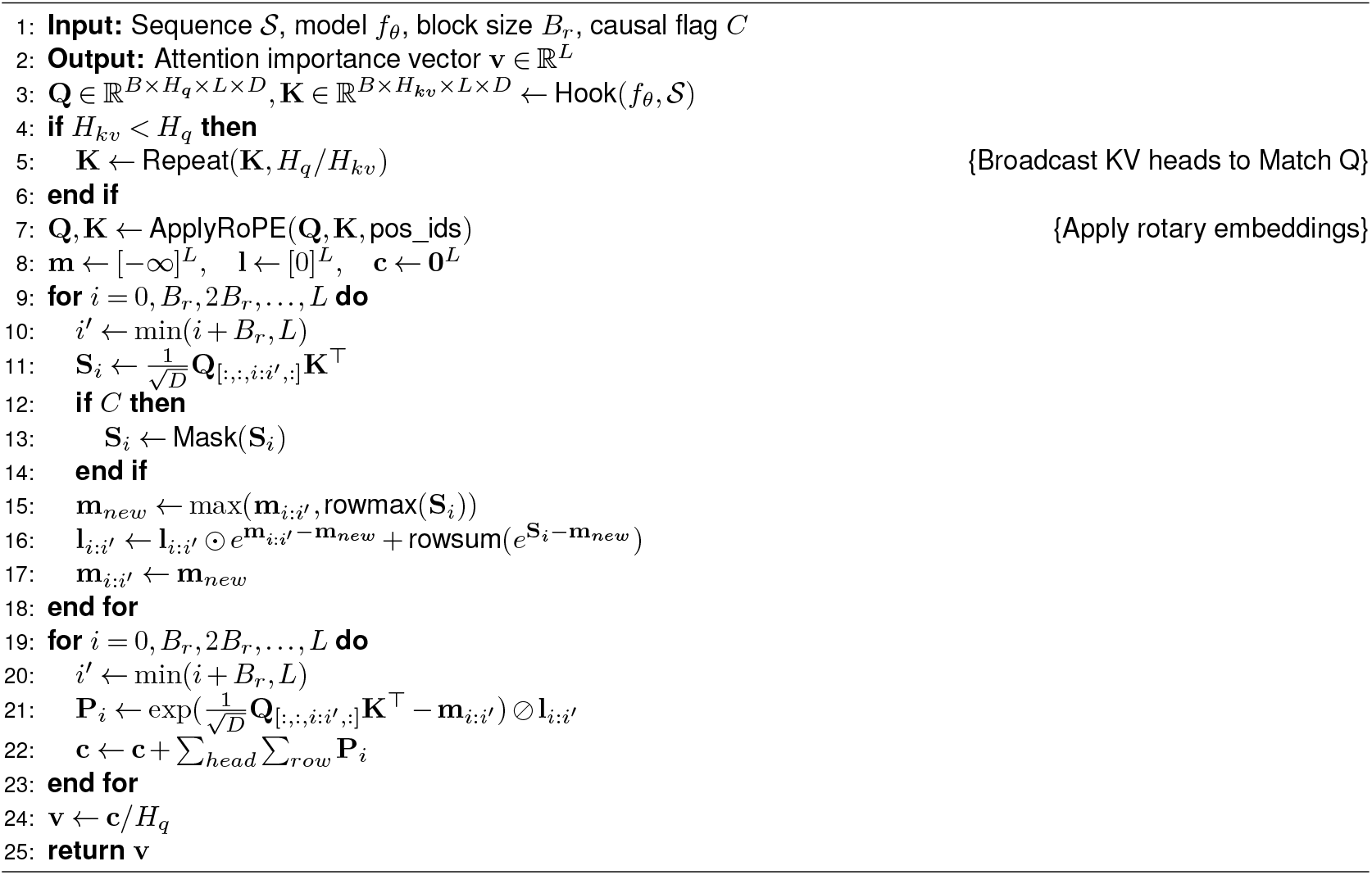

### B.2 Dynamic Coordinate Alignment

Post-extraction, the vector **v** corresponds to token indices. To map these to genomic coordinates *p*, we apply the following rules to handle variants (Indels):

- **Insertions (Shared Coordinates):** Tokens generated by an insertion share the genomic coordinate of the preceding anchor base. The attention score for coordinate *p* is calculated as the aggregation of all tokens mapped to *p*.
- **Deletions (Placeholder Encoding):** Deleted regions retain their genomic keys in the coordinate map but generate no tokens. This ensures that the relative distance of downstream tokens remains biologically consistent.

### C. Long-Sequence Strategy and Hyperparameter Ablation

#### C.1 Empirical Benchmarking on NVIDIA H100

The strategy transition thresholds (4k and 128k) were established through rigorous benchmarking on a single **NVIDIA H100 (80GB) GPU**.

##### Scalability Constraints

As detailed in Table 5, **Vanilla Attention** memory usage grows quadratically. While efficient for short contexts, it reaches 53.7 GB at *L* = 8, 192 and triggers Out-of-Memory (OOM) errors at 32k. Conversely, **FlashAttention** (Table 6) maintains linear memory scaling, enabling processing up to 131, 072 (128k) bp. However, at 128k, the latency per sequence rises to 15.47s, and memory usage hits 42.9 GB. To ensure robust throughput and prevent instability at extreme lengths, we define *L* = 128k as the upper bound for exact global attention, switching to chunked processing thereafter.

**Table 5.** Computational Cost of Vanilla Attention (Single H100)

| # GPUs | Length ($L$) | Latency (s) | Memory (MB) |
| --- | --- | --- | --- |
| 1 | 2,048 | 0.2359 | 21,967 |
| 1 | 4,096 | 0.4221 | 28,425 |
| 1 | 8,192 | 1.1600 | 53,677 |
| 1 | 32,768 | OOM | OOM |

**Table 6.**
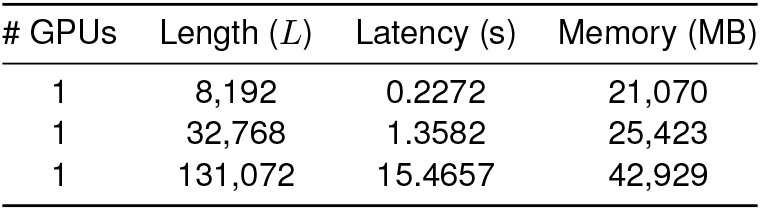
Computational Cost of FlashAttention (Single H100)

| # GPUs | Length ($L$) | Latency (s) | Memory (MB) |
| --- | --- | --- | --- |
| 1 | 8,192 | 0.2272 | 21,070 |
| 1 | 32,768 | 1.3582 | 25,423 |
| 1 | 131,072 | 15.4657 | 42,929 |

#### C.2 Chunking Hyperparameter Optimization

For the chunking strategy (*L >* 128k), we performed an ablation study to select the optimal window size *C* and overlap *O*. We measured the Area Under the Precision-Recall Curve (AUPRC) as a proxy for the model’s Signal-to-Noise Ratio (SNR) in detecting regulatory variants.

As shown in Figure S1, we evaluated chunk sizes ranging from 4k to 32k. The configuration of *C* = 8, 192 (Chunk Size) and *O* = 4, 096 (Overlap) yielded the highest AUPRC across multiple sequence lengths (green line). While larger chunks (32k) theoretically offer more context, our analysis suggests they introduce excessive background noise that dilutes the local signal, in addition to higher computational overhead (see Figure S2). Conversely, smaller chunks (4k) fracture long-range dependencies essential for distal regulation. Thus, the 8k/4k setup provides the optimal trade-off between signal fidelity and computational cost.

## Supplementary Note 2: Data & Experiments Details

### A. Real Dataset Characteristics

#### A.1 Sample Classification and Ground Truth

We designed this real-world evaluation to assess whether ATLAS can recover known related loci under realistic population heterogeneity and identify plausible disease-associated signals beyond curated annotations.

The *β*-thalassemia dataset was retrieved from a cross-sectional whole-genome sequencing study investigating the clinical heterogeneity of hemoglobinopathies. The cohort comprises 1,429 individuals, classified into three groups based on clinical severity:

- **Carriers (***N* = 409): Individuals who generally exhibit no overt clinical symptoms, although mild anemia and microcytosis may be observed in hematological tests.
- **Mild Cases (Thalassemia Intermedia, TI**, *N* = 245): Classified as non-transfusion dependent thalassemia (NTDT), characterized by low or intermittent transfusion dependence and relatively milder clinical manifestations.
- **Severe Cases (Thalassemia Major, TM**, *N* = 775): Classified as transfusion-dependent thalassemia (TDT),characterized by high transfusion dependence and severe disease progression.

**Figure S1.**
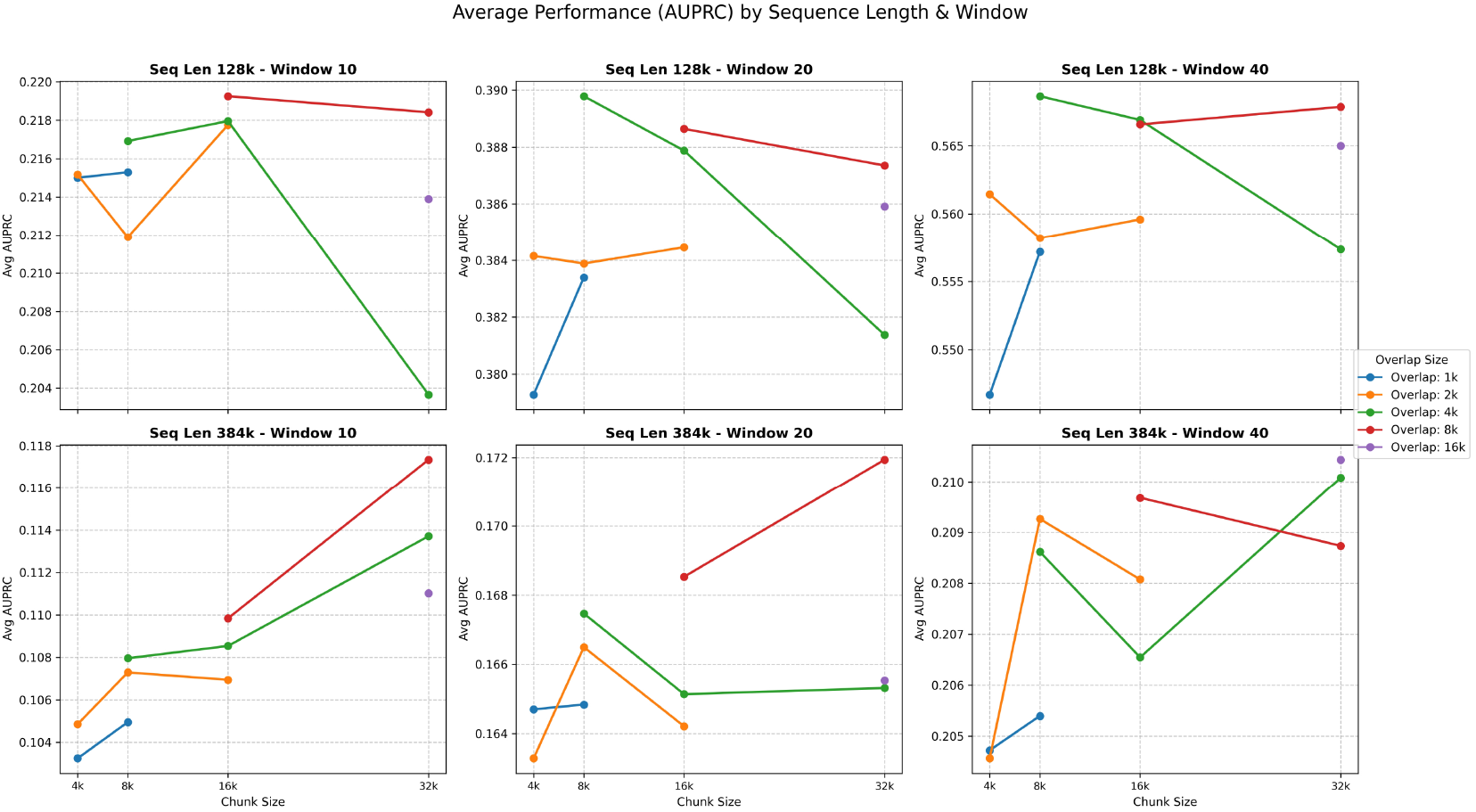
Ablation Study - Performance Analysis (AUPRC). Comparison of variant detection accuracy across varying chunk sizes and overlaps. The 8k chunk size with 4k overlap (green line) consistently shows the optimal trade-off between signal detection and context capture.

**Figure S2.**
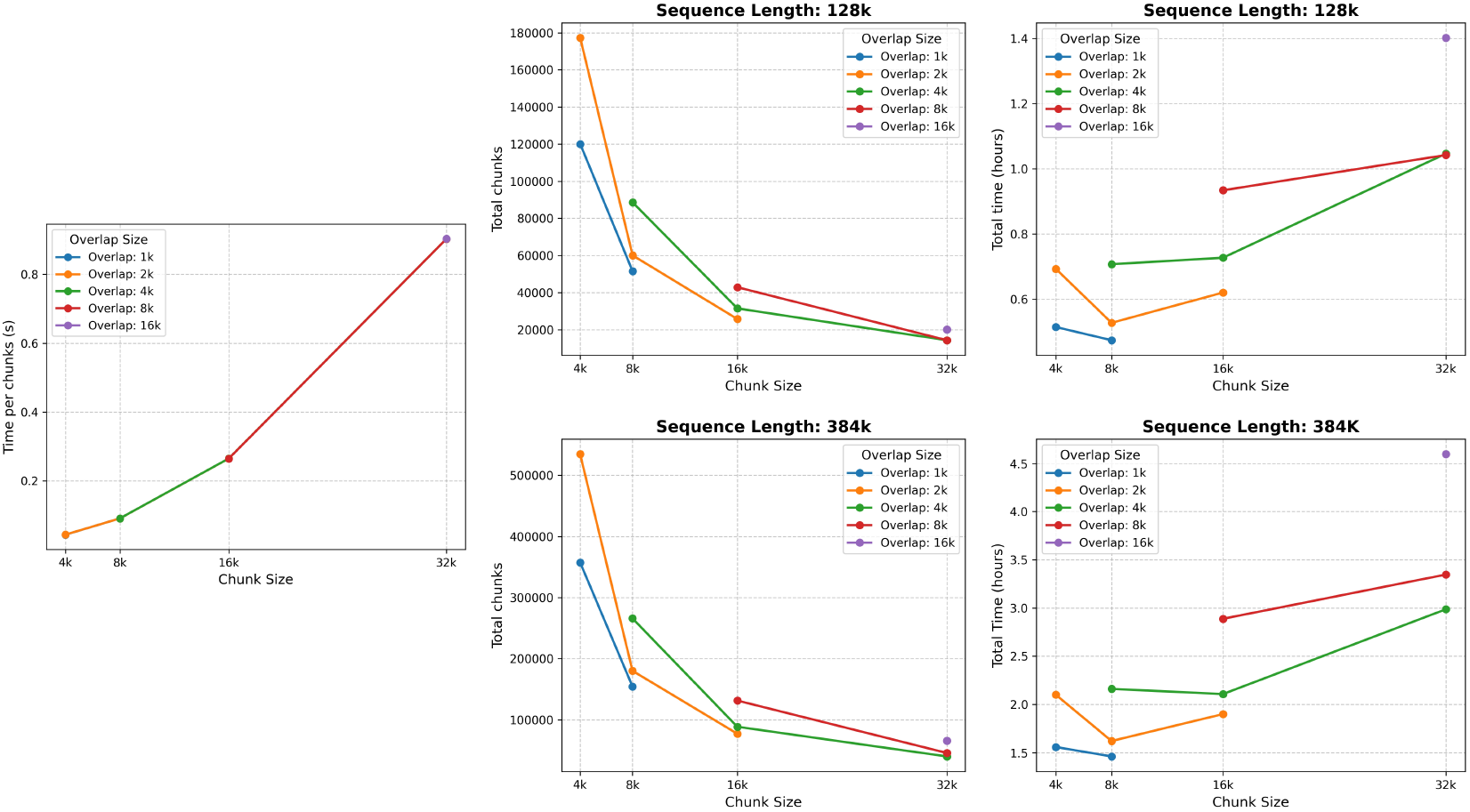
Ablation Study - Computational Cost Analysis. Total processing time (hours) and chunk generation overhead. While larger chunks (32k) reduce total time, they degrade detection performance (as shown in Figure. S1). The 8k size maintains a reasonable computational cost.

The clinical classification of these samples serves as the ground truth labels for all subsequent analyses.

### A.2 Genomic Scope

We analyzed all protein-coding genes located on chromosome 11 with lengths shorter than 512 kb. This comprises a total of **1**,**031 genes**. The length distribution is as follows:

- *<* 64 kb: 1,049 genes (Note: strictly adhering to the *<*512kb filter).
- 64 kb − 128 kb: 145 genes.
- 128 kb − 256 kb: 72 genes.
- 256 kb − 512 kb: 35 genes.

### B. Data Processing Pipeline

#### B.1 Phasing and Haplotype Construction

Haplotype phasing was performed on the original VCF files using **Beagle v4**. Default parameters were applied with genotype probability output enabled (`gp=true`) and a sliding window size of`window=10.0.` This procedure phased unphased genotypes into two haplotypes (`hap1 and hap2`) for each sample.

#### B.2 Sequence Construction and Coordinate Mapping (VCF2CSV)

We constructed haplotype-specific sequences by mapping VCF variants to the GRCh38 reference genome. For each sample, sequences were processed from the 5’ end to the 3’ end to handle coordinate shifts dynamically. The mapping rules are defined as follows:

1. **No Variant:** The reference base is retained, and its absolute genomic position is recorded.
2. **SNPs:** The reference base is replaced by the alternative allele, recording the original reference position.
3. **Deletions:** Bases are excluded from the sequence, and their positions are *not* recorded (only remaining bases retain position tags).
4. **Insertions:** Inserted bases are included sequentially. Crucially, the position of the preceding reference base is **repeated** for each inserted base to maintain alignment with the reference coordinate system.

##### Strand Handling

For genes located on the negative strand, the generated sequences were reverse-complemented, and the corresponding position arrays were reversed to maintain alignment with the reference genome coordinate order.

### C. Synthetic Dataset Generation

We provide the detailed statistics of the synthetic datasets constructed for robustness evaluation in Table 7. The synthetic generation process involved injecting variants into intergenic regions derived from chromosome 11.

**Table 7.** Statistics of the Synthetic Datasets across different sequence lengths.

| Seq Length | Carriers | Mild | Severe | Target Variants (SNP/Indel) | Target Prop. |
| --- | --- | --- | --- | --- | --- |
| 4 kb | 409 | 245 | 775 | 8/0 | 100% |
| 20 kb | 409 | 245 | 775 | 8/2 | 100% |
| 128 kb | 409 | 245 | 775 | 15/5 | 100% |
| 384 kb | 409 | 245 | 775 | 15/5 | 100% |

### D. Baseline Settings

#### D.1 Foundation Model

We utilize **Genos-10b-v1** as the foundation model baseline. The input sequences are padded or truncated to match the model’s maximum context length where necessary.

#### D.2 GWAS Configuration

Genome-Wide Association Studies (GWAS) are conducted using **PLINK2**. We apply a unified threshold strategy for fair comparison.

- **Quality Control (QC):** Variants with missingness *>* 1% (`–geno 0.01`) and Minor Allele Frequency *<* 1% (`–maf 0.01`) were removed.
- **Locus Identification:** A GWAS locus is counted as co-identified if its position falls within an attention-derived cluster interval defined as:

**Listing 1.**
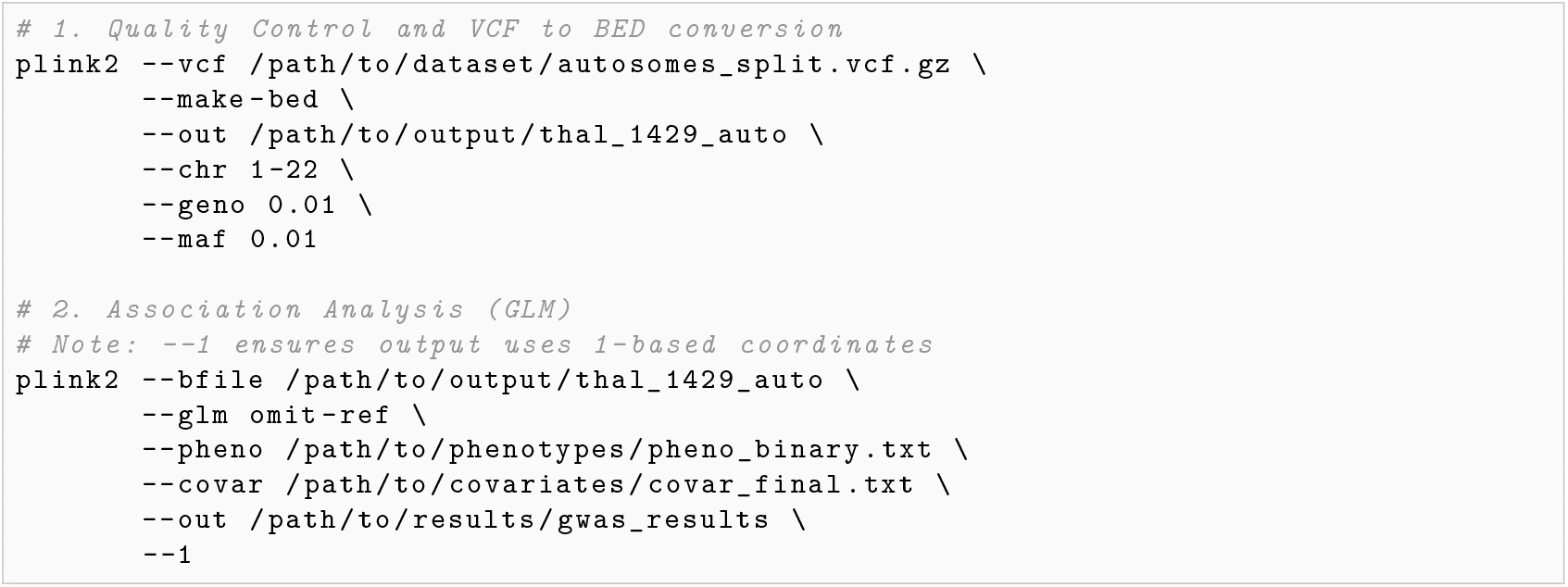
GWAS Pipeline Commands (PLINK2)

– Forward strand: [*start* − 10, *end*]
– Reverse strand: [*start, end* + 10]

The specific PLINK2 commands used for Quality Control (QC) and Association Analysis are listed below:

## Supplementary Note 3: Baseline Model Configurations

To benchmark our approach, we compare it against state-of-the-art genomic language models in different pre-train scopes to verify the influences of model sizes and training data on downstream tasks. Table 8 summarizes the architectures and specifications. Notably, all selected models utilize a **single-nucleotide tokenizer**, ensuring a fair comparison at the base resolution level. Regarding the Evo 2 series, given its hybrid StripedHyena architecture where attention heads are interspersed with convolution operators, we identified the best-performing layers reported in the original study and selected the nearest attention layer immediately preceding them for analysis.

**Table 8.** Summary of Baseline Models. The “Extraction Layer” column indicates which layer’s attention weights were used for analysis.

| Model Name | Architecture | Pre-train Scope | Data Volume | Context Length | Param Size | Extraction Layer |
| --- | --- | --- | --- | --- | --- | --- |
| **Evo2-1B-Base** | StripedHyena 2 | Multispecies | 1T bp | 8K bp | 1.1B | Layer 10 |
| **Evo2-7B-Base** | StripedHyena 2 | Multispecies | 2.1T bp | 8K bp | 6.5B | Layer 24 |
| **Evo2-40B** | StripedHyena 2 | Multispecies | 9.3T bp | 1M bp | 40B | Layer 17 |
| **Genos-1.2B** | MoE (Mixture of Experts) | Human | 1600B tokens | 1M | 1.2B | Last Layer |
| **Genos-10B** | MoE (Mixture of Experts) | Human | 2200B tokens | 1M | 10B | Last Layer |
| **Genos-10B-v2** | MoE (Mixture of Experts) | Human | 6286B tokens | 1M | 10B | Last Layer |
| **LucaOne** | Transformer | Multispecies | 36.95B bp | 1k | 1.8B | Last Layer |
| **LucaVirus** | Transformer | Viral sequences | 50B bp | 3k | 1B | Last Layer |

## Supplementary Note 4: Evaluation Metrics for Signals

### A. Overview

To evaluate the spatial concentration of differential signals around true variant positions, we develop a set of evaluation metrics. These metrics quantify how well the predicted signals localize to regions surrounding known variants. To enable fair comparison across sequences of different lengths (e.g., 4 kb vs. 384 kb), we additionally introduce length-normalized variants of key metrics.

### B. Notation

Let **s** = (*s*_1_, *s*_2_, …, *s*_*N*_) denote the signal scores (absolute log_2_ fold-change of attention) at *N* genomic positions, where *p*_*i*_ denotes the genomic coordinate of position *i*. Given a set of *K* true variant positions 𝒱 = {*v*_1_, *v*_2_, …, *v*_*K*_}, we define a binary label *y*_*i*_ ∈ {0, 1} indicating whether position *i* falls within a window around any variant:

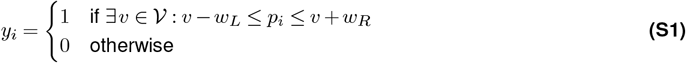

where *w*_*L*_ and *w*_*R*_ are the left and right window boundaries, respectively (tested at 3 bp, 5 bp, 10 bp, and 20 bp). We further define 𝒯 = {*i* : *y*_*i*_ = 1} as the set of positions within true variant windows (positives), ℱ = {*i* : *y*_*i*_ = 0} as the set of background positions (negatives), with *n*_*T*_ = | 𝒯 | and *n*_*F*_ = | ℱ | denoting their respective counts. For spatial metrics, we define *d*_*i*_ = min_*v*∈ 𝒱_ |*p*_*i*_ − *v*| as the distance from position *i* to the nearest variant.

### C. Primary Metric: AUPRC

We adopt the **Area Under the Precision–Recall Curve (AUPRC)** as the primary metric to quantify ranking performance under severe class imbalance. AUPRC summarizes the precision-recall trade-off across all classification thresholds:

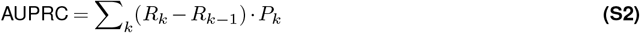

where *P*_*k*_ and *R*_*k*_ are the precision and recall at the *k*-th threshold. AUPRC is particularly informative when positive cases are rare. The baseline (random classifier) AUPRC equals the proportion of positives (*n*_*T*_ */N*); thus, AUPRC values should be interpreted relative to this baseline when comparing across sequences of different lengths. As shown in Figure S7, we evaluated performance across varying window sizes to ensure robustness.

### D. Complementary Metrics

To overcome the limitations of threshold-based metrics, we introduce complementary indicators measuring magnitude contrast, signal efficiency, and spatial precision.

#### D.1 Signal-to-Noise Ratio (SNR)

Categorized as *Magnitude Contrast*, SNR measures how biologically distinct the variant signal is from the background noise floor:

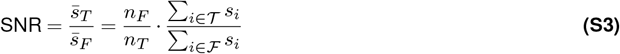

Where 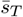 and 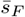 denote the mean signal in true variant windows and background regions, respectively. SNR *>* 1 indicates that signals are, on average, stronger near variants than in background regions; higher values indicate better signal specificity. Figure S8 illustrates the SNR distribution.

#### D.3 Fraction of Signal in Windows (FRiW)

Categorized as *Signal Efficiency*, this metric is analogous to the FRiP score in ChIP-seq. It quantifies the proportion of total signal that falls within true variant windows:

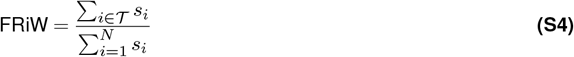

A low FRiW implies that despite a potentially high AUPRC, the majority of the model’s attention mass is allocated to regions outside variant windows (see Figure S9).

### D.3 Signal-Weighted Mean Distance

Categorized as *Spatial Precision (Threshold-Free)*, this metric measures the average distance of signal from the nearest variant, weighted by signal intensity, removing the need for hard window boundaries:

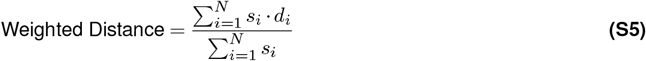

A **lower score** is better, indicating that the attention mass is concentrated physically closer to the causal variants (see Figure S10).

### E. Length-normalized Metrics

To enable fair comparison across sequences of vastly different lengths, we introduce length-normalized variants of the above metrics. These normalized metrics account for the expected baseline values under uniform signal distribution, making them suitable for cross-scale comparisons.

#### E.1 FRiW Enrichment

To account for varying sequence lengths and window sizes, we normalize FRiW by the expected fraction under a uniform signal distribution:

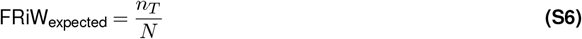

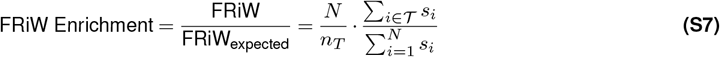

This fold-enrichment metric is length-independent. A value of 1 implies a signal distribution equivalent to random expectation (uniform background); values *>* 1 indicate a significant enrichment of attention mass within variant windows.

#### E.2 Normalized Weighted Distance

To enable comparison across sequences of different lengths, we normalize the weighted distance by the characteristic length scale of the sequence. Under a uniform distribution assumption with *K* variants, the expected distance to the nearest neighbor scales linearly with *L/K*. We therefore define:

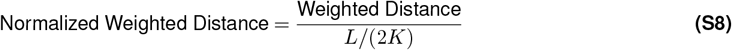

This normalization factor *L/*(2*K*) serves as a first-order approximation of the expected random distance, facilitating fair comparison of spatial accuracy across varying genomic scales.

#### E.3 Mean Percentile Rank of True Positions

This rank-based metric evaluates where true variant positions fall in the ranked signal distribution:

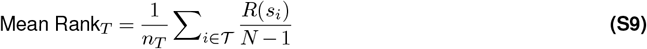

where *R*(*s*_*i*_) is the rank of *s*_*i*_ among all scores (0 for lowest, *N*− 1 for highest). This metric ranges from 0 to 1, where 0.5 indicates performance equivalent to random ranking, and values approaching 1 indicate that positions near variants consistently have high signal scores. As a rank-based metric, Mean Percentile Rank is completely length-independent.

## Supplementary Note 5: Gene-Level Statistical Descriptors

The gene-level analysis aims to identify distribution differences of attention scores across entire genes. For a gene with *M* positions and attention scores **s** = (*s*_1_, *s*_2_, …, *s*_*M*_), we compute **17 descriptors** categorized into four groups. In our experiments, **Max, Std** (*σ*), **Top5%Mean, CV, Median, IQR**, and **Entropy** provide the most significant separation between the informative (HBB) and control genes, suggesting that both the magnitude of extreme values and the overall distribution shape are informative for distinguishing regulatory patterns.

### A. Location/Scale (10 metrics)

- **Mean:** The arithmetic mean of all scores:

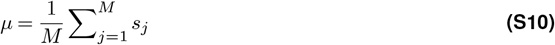
- **Median:** The middle value when scores are sorted, i.e., *Q*_0.50_.
- **Top5% Mean:** Mean of the top 5% highest scores:

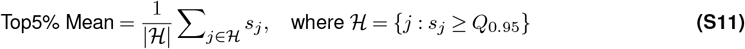
- **Low5% Mean:** Mean of the bottom 5% lowest scores:

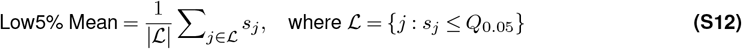
- **Max:** The maximum score across all positions:

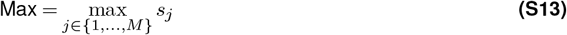
- **Standard Deviation:** Measures the spread of scores around the mean:

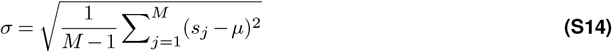
- **Coefficient of Variation (CV):** Scale-normalized dispersion:

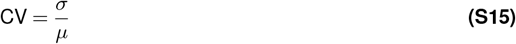
- **Interquartile Range (IQR):** The range between the 25th and 75th percentiles:

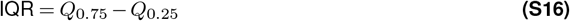
- **10th Percentile** (*Q*_0.10_) and **90th Percentile** (*Q*_0.90_): Values below which 10% and 90% of scores fall, respectively.

### B. Distribution Shape (3 metrics)

- **Skewness:** The standardized 3rd central moment, measuring distribution asymmetry:

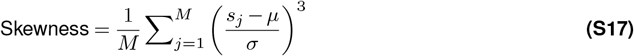

Positive skewness indicates a right-tailed distribution; negative skewness indicates a left-tailed distribution.
- **Kurtosis:** The standardized 4th central moment, measuring tail heaviness:

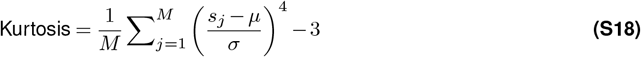

Values *>* 0 (leptokurtic) indicate heavy tails; values *<* 0 (platykurtic) indicate light tails.
- **Mode:** The most frequent value, computes via a fixed-bin histogram as the center of the bin with maximum count.

### C. Peak Structure (3 metrics)

To capture local regulatory motifs and identify regions of concentrated attention:

- **Peak Count:** Number of local maxima identified, where position *j* is a local maximum if *s*_*j*_ *> s*_*j*−1_ and *s*_*j*_ *> s*_*j*+1_.
- **Peak Density:** PeakCount normalized by gene length:

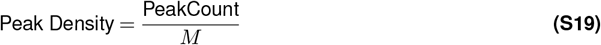
- **Peak Mean:** Mean attention score at peak summits:

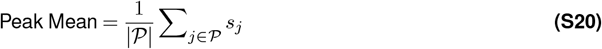

where 𝒫 is the set of positions identified as local maxima.

### D. Information (1 metric)

- **Shannon Entropy:** Measures the sparsity or concentration of the attention distribution. After normalizing scores to a probability distribution 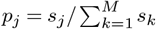 the entropy is computed as:

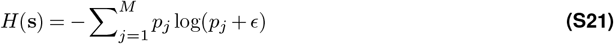

where *ϵ* is a small constant for numerical stability. Lower entropy indicates more concentrated (sparse) attention; higher entropy indicates more uniform distribution.

**Figure S3.**
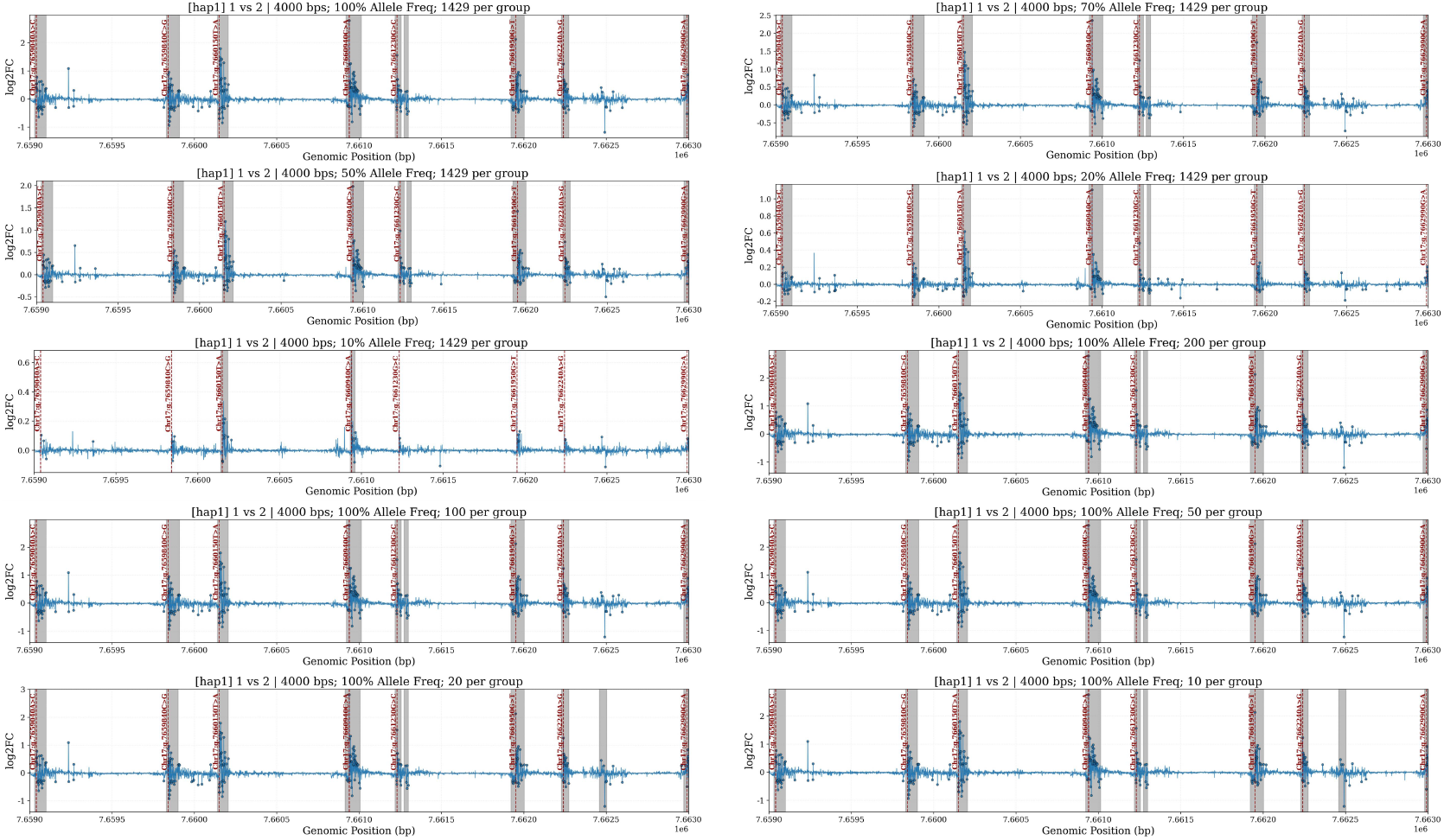
ATLAS results under different allele frequencies and cohort sizes. Red vertical lines are the positions of the synthetic variants. Dots are the bases with significant attention differences. The gray areas are the clusters.

**Figure S4.**
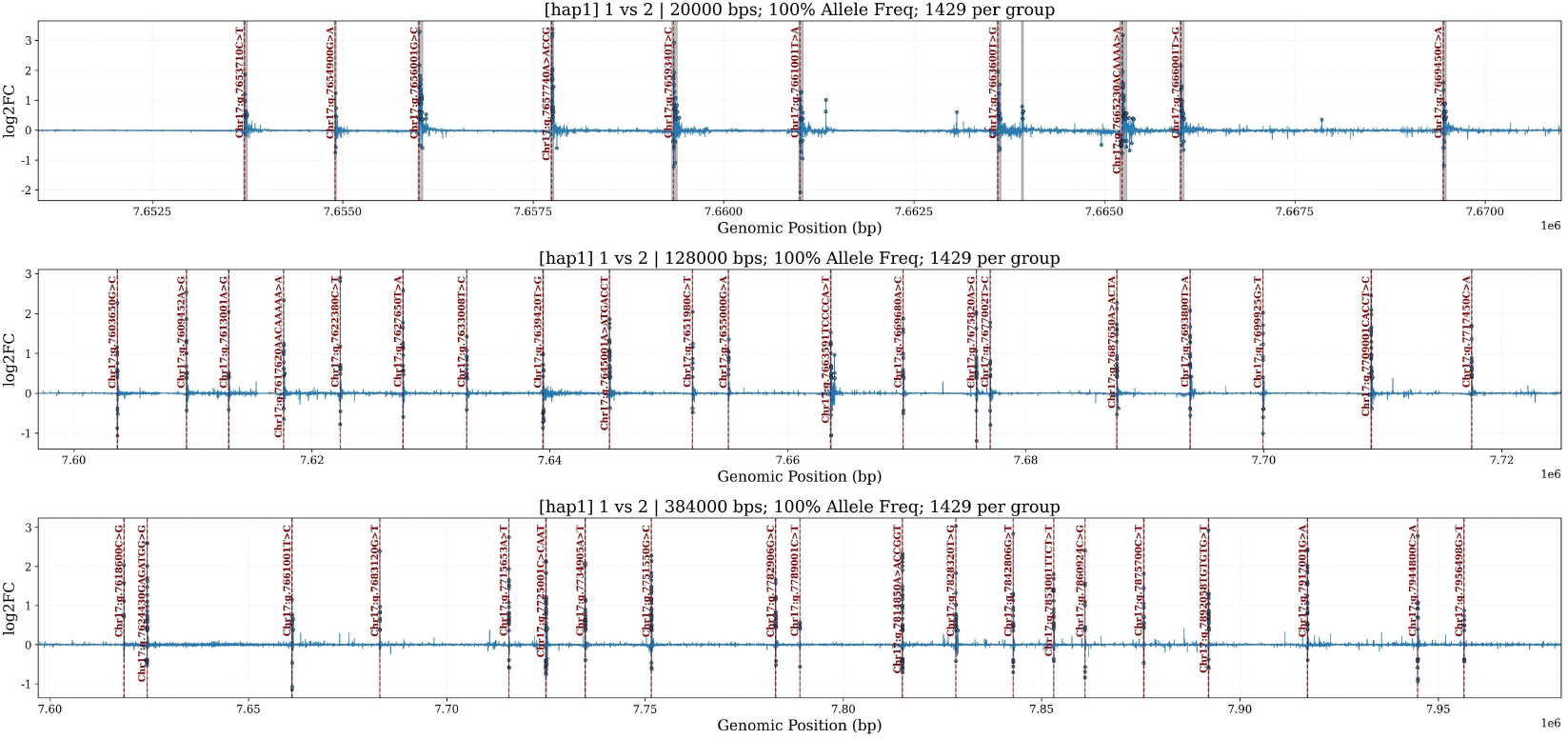
ATLAS results under different sequence lengths. Notations are the same as Figure S3. The gray areas may not be clearly visible in long sequences.

**Figure S5.**
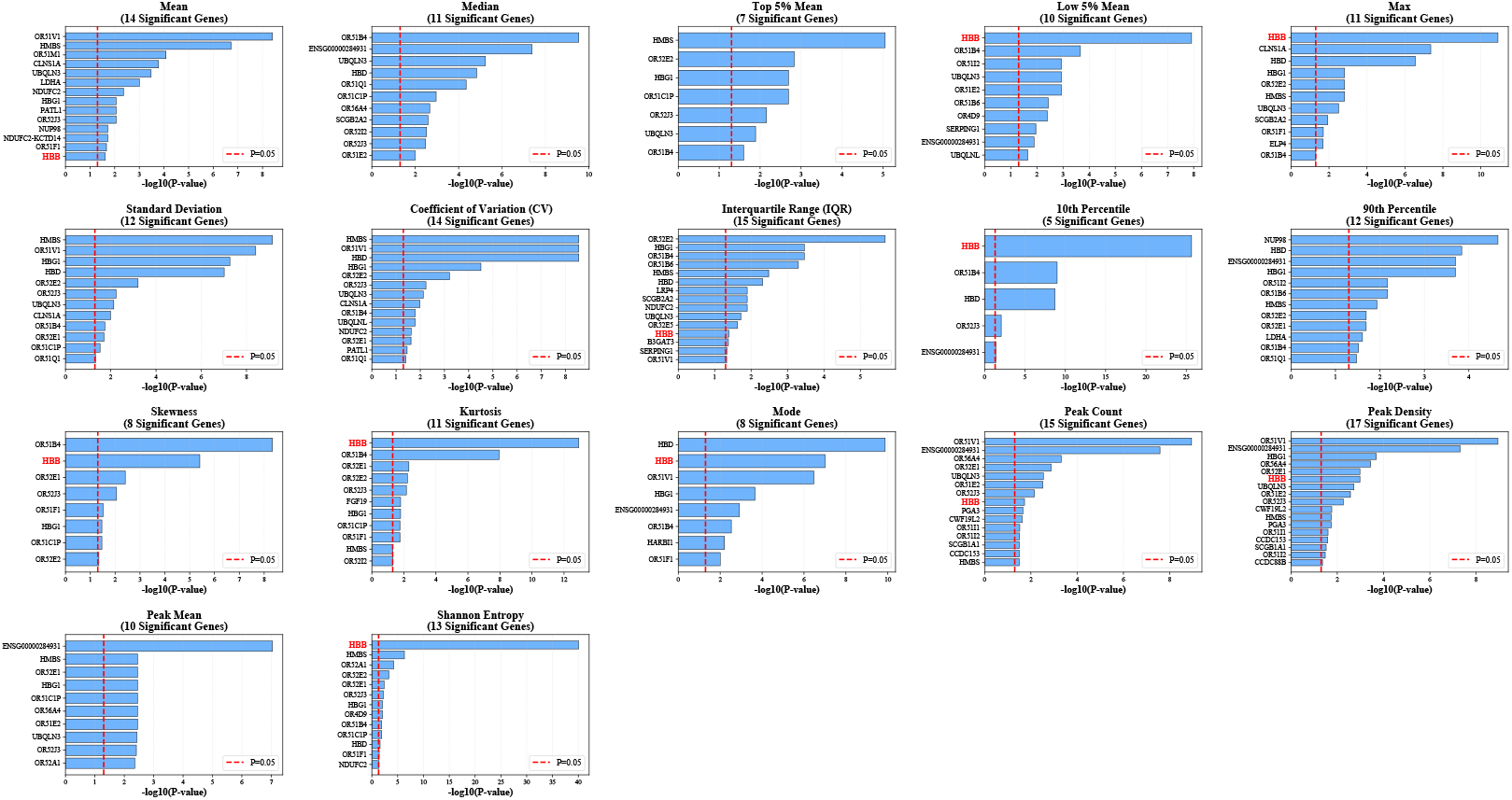
The significant genes selected by the 17 distribution descriptor in the gene-level analysis. The results are retrieved from the haplotype 1 attention scores

**Figure S6.**
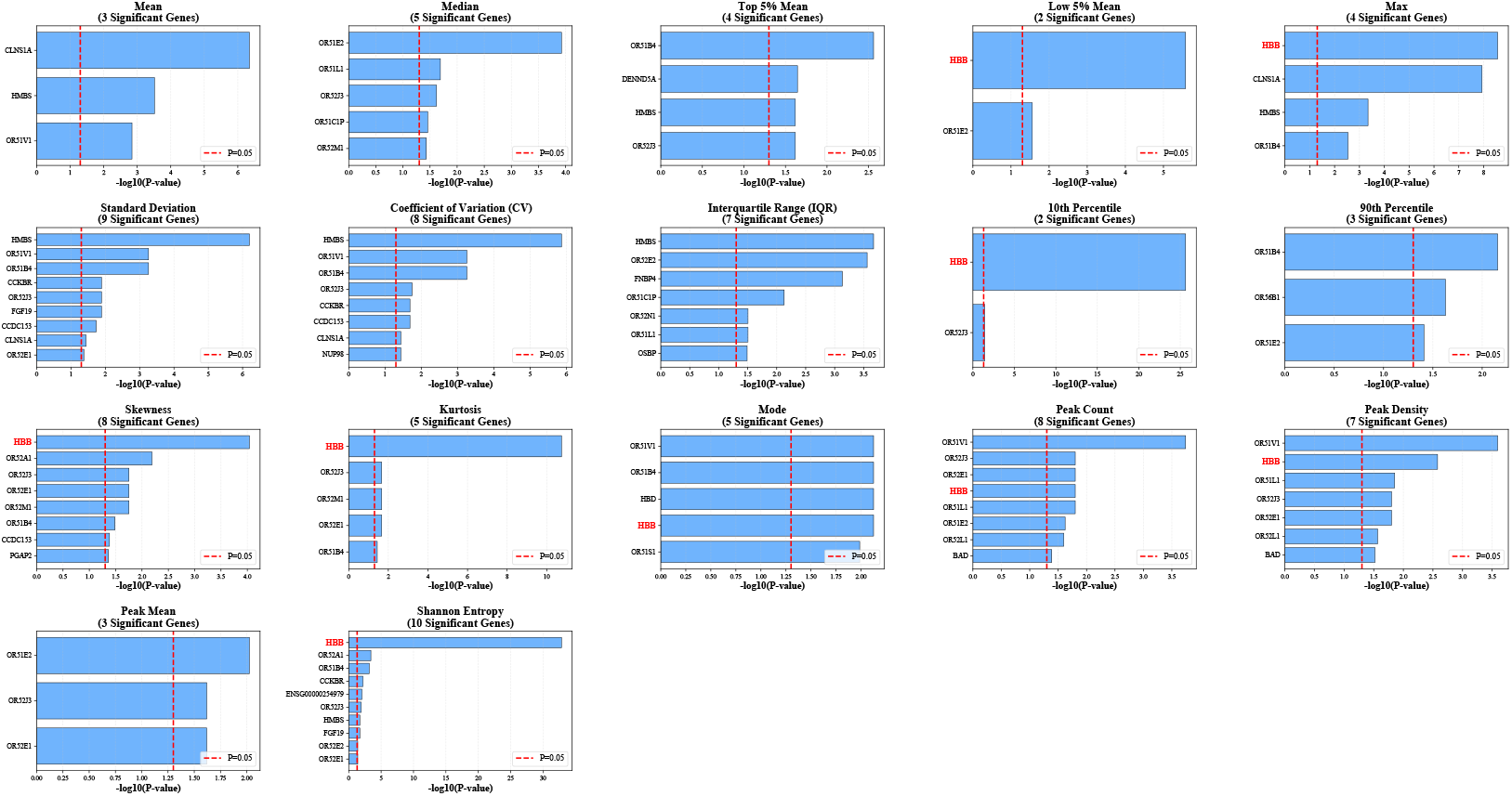
The significant genes selected by the 17 distribution descriptor in the gene-level analysis. The results are retrieved from the haplotype 2 attention scores

**Figure S7.**
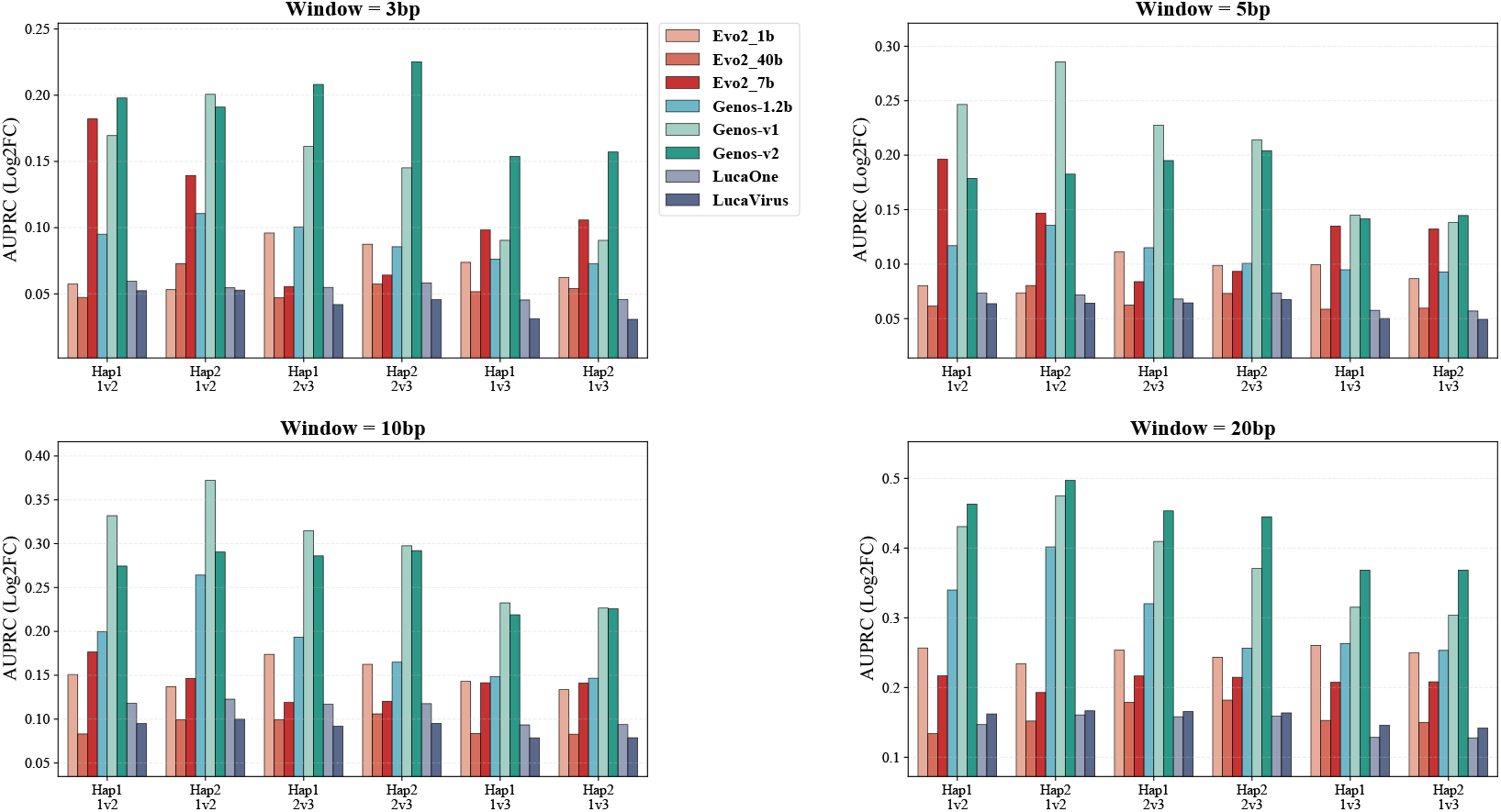
Evaluation of Detection Quality - AUPRC Analysis (Higher is better). Evaluation of ranking performance across different window sizes (3bp, 5bp, 10bp, 20bp). While AUPRC measures the ranking order, the Genos-v2 model (brown) consistently demonstrates strong ranking capabilities across most haplotype comparisons.

**Figure S8.**
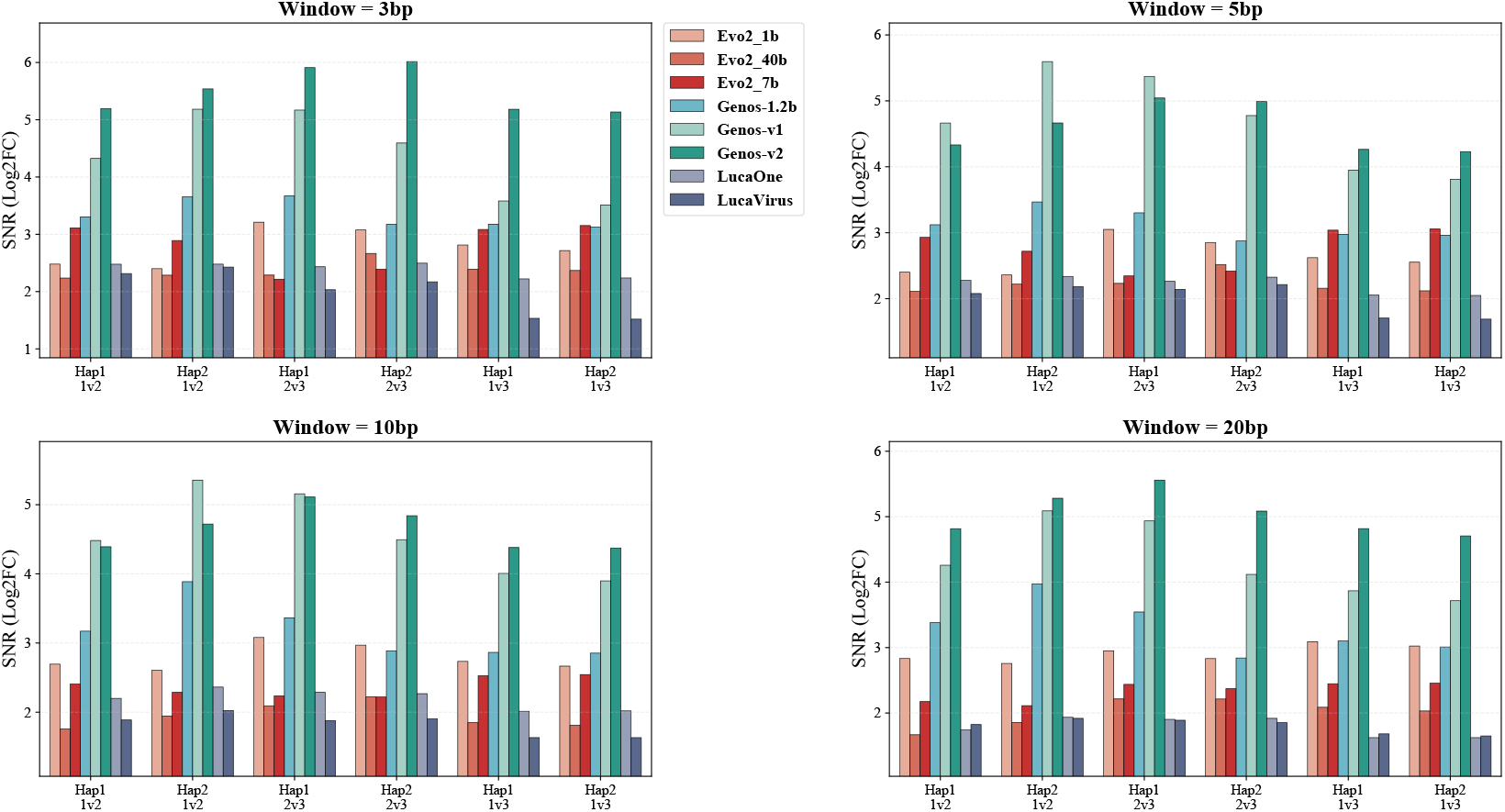
Evaluation of Detection Quality - Signal-to-Noise Ratio (SNR) Analysis (Higher is better). Assessing the magnitude contrast between variant signals and background noise. A higher SNR confirms that the identified signals have a significant magnitude difference compared to the background, indicating that the model’s attention peaks at **risk variants** are biologically distinct from the noise floor.

**Figure S9.**
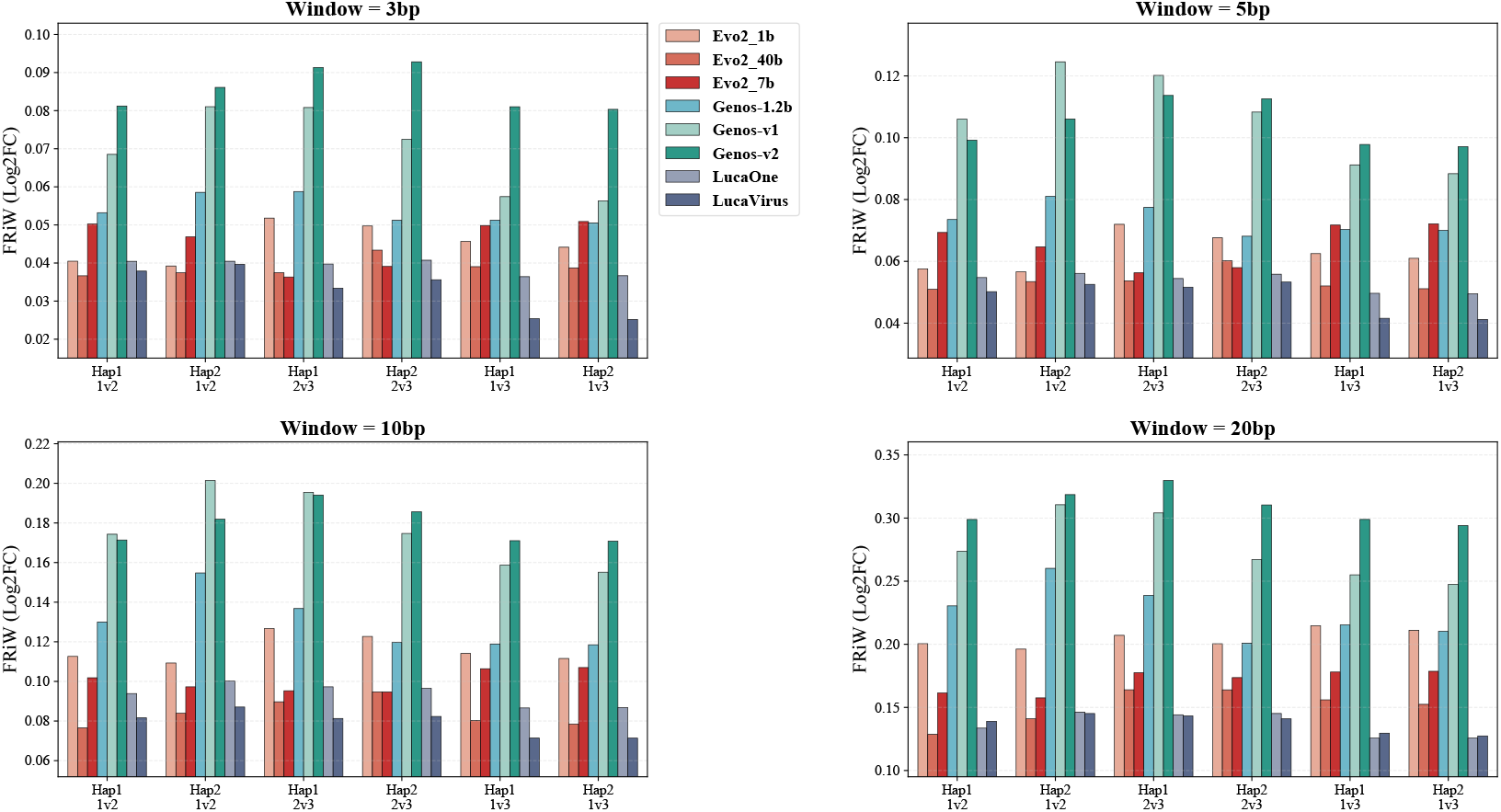
Evaluation of Attention Efficiency - Fraction of Signal in Windows (FRiW) (Higher is better). Quantifying the global attention budget allocation. Models with higher FRiW scores are more efficient, concentrating their attention mass into the relevant variant windows rather than dispersing it across the sequence.

**Figure S10.**
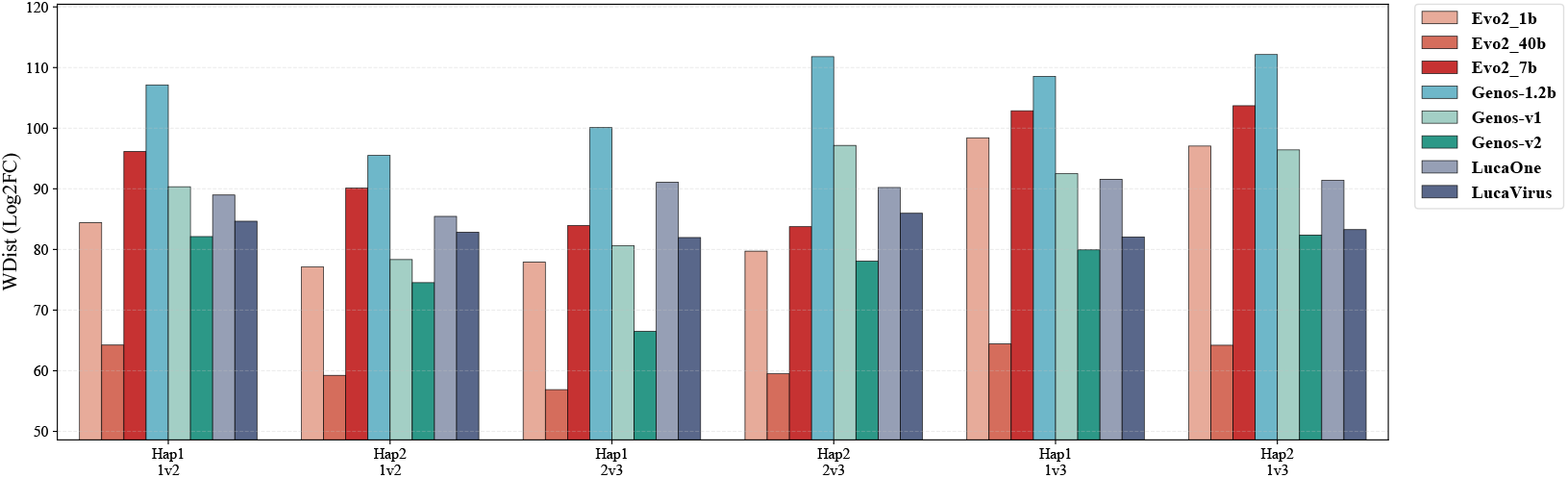
Evaluation of Spatial Precision - Distance-Weighted Mean (Lower is better). A threshold-free metric measuring spatial precision. Lower values indicate that high attention scores are physically closer to the **target variants**, minimizing spatial deviation.

## Notes

### Competing Interest Statement

The authors have declared no competing interest.

